# Susceptibility-induced internal gradients reveal axon morphology and cause anisotropic effects in the dMRI signal

**DOI:** 10.1101/2023.05.01.538981

**Authors:** S. Winther, H. Lundell, J. Rafael-Patiño, M. Andersson, J-P. Thiran, T. B. Dyrby

## Abstract

Diffusion-weighted MRI is our most promising method for estimating microscopic tissue morphology in vivo. The signal acquisition is based on scanner-generated *ex*ternal magnetic gradients. However, it will also be affected by susceptibility-induced *in*ternal magnetic gradients caused by interaction between the tissue and the static magnetic field of the scanner. With 3D in silico experiments, we show how internal gradients cause morphology-, compartment-, and orientation-dependence of spin-echo and pulsed-gradient spin-echo experiments in myelinated axons. These effects are unseen in previous 2D modelling. For an ex vivo monkey brain, we observe the orientation-dependency generated only when including non-circular cross-sections in the in silico morphological configurations, and find orientation-dependent deviation of up to 17% for diffusion tensor metrics. Our findings underline the importance of accounting for realistic 3D axon morphology in modelling. Interestingly, the morphology-specific orientation-dependency trends show potential for a novel sensitivity to morphology, which is not attainable by the theoretical diffusion-weighted MRI signal itself.

## 1 Introduction

Diffusion-weighted MRI (dMRI) can reveal microscopic tissue morphology of the brain by measuring water diffusion properties within the tissue. The image contrast is induced by manipulating the phase coherence of nuclear spins with external magnetic gradients applied with the MR scanner. However, because different tissue components possess different magnetic susceptibilities, internal magnetic gradients are induced in the tissue and will contribute to the image contrast. The internal gradients are morphology- and orientation-dependent, and correlate with the strength of the MR scanner’s **B_0_**-field; thereby having a larger impact in the emerging ultra-high field scanners. While some MRI sequences (e.g. gradient-echo-based sequences) are designed to target such internal gradients, their contributions to the signal are commonly disregarded in dMRI, and may hence bias the signal interpretation. However, if the contributions are accounted for in the signal modelling, the very same internal gradients may carry a novel insight into the underlying microstructure. The most commonly applied sequence in dMRI is the pulsed-gradient spin-echo (PGSE) [1]. In the presence of internal gradients and diffusion beyond a characteristic length, the isolated spin-echo (SE) will be attenuated due to dephasing [2]. This is measured as a T2 effect. Moreover, because cross-terms arise between the external pulsed gradients and the internal gradients [1][3], the PGSE signal becomes sensitive not only to diffusion in specific directions (as desired), but also to internal gradients in specific directions. These aspects may cause both the SE and PGSE signals from identical microstructure to deviate if oriented differently w.r.t. **B_0_**.

Diffusion-weighting of the PGSE sequence can be signified by a *b*-vector. The desired b-vector *b_des_* is given from the user-specified scan sequence parameters. However, in the presence of internal gradients the effective b-vector *b_eff_* is a sum of contributions also from the internal gradients *b_int_*, and from cross-terms between the internal and external gradients *b_cross_* [1][3]. This means that *b_eff_* will depend on the internal gradients of the given tissue, and thereby may reflect the morphology of the tissue. Furthermore, because the impact of the internal gradients may be different across tissue compartments, the common normalization w.r.t. *S*(*b_des_* = 0) may shift the weighting of different compartments.

Different strategies have been developed to correct for cross-terms in PGSE sequences. Cross-terms with internal gradients that appear linear over a characteristic diffusion length can be corrected by e.g. taking the geometric average of the signals from two b-vectors of opposite polarity [4][5], using bipolar pulsed gradients on each side of the inversion pulse [6], or numerical optimization scheme for general waveforms [3]. However, to cancel out cross-terms with internal gradients that appear nonlinear, more advanced sequences are required. A 13-interval STEAM sequence [7] has proven capable - but is only practically applicable at long diffusion times. Effects of internal gradients in PGSE must therefore be addressed as part of the signal modelling. To do so, we need a better understanding of the mechanisms behind these internal gradients and their effects on dMRI signals.

In brain white matter, the main contribution to internal gradients is believed to come from myelin [8][9]. Myelinated axons have commonly been modelled as infinitely long non-touching straight cylinders [10]. For the special case of an infinitely long isolated straight coaxial cylinder, the susceptibility-induced gradients have been solved analytically; both when considering the susceptibility of myelin as isotropic [11], and as anisotropic (i.e. modelling the polarity of the lipids within the myelin) [12][13]. In either case, gradients are induced only in the extra-axonal and myelin compartments, while the intra-axonal compartment is affected homogeneously; thereby leaving any intra-axonal SE signal unaffected. Furthermore, gradients are induced only when the orientation of the cylinder is non-parallel with the applied magnetic field.

From recent 3D synchrotron X-ray nano-holotomography (XNH) in monkey brain [14], and 3D electron microscopy in rodent brains [15][16], the morphology of myelinated axons has been found to demonstrate non-circular cross-sections, longitudinal diameter variations, and tortuous trajectories. Tortuous trajectories have previously been quantified by micro-dispersion [14]. These deviations from straight coaxial cylinders may induce internal gradients also in the intra-axonal compartment which have not been accounted for previously. More realistic axon morphology has been simulated in 2D based on ellipses and segmentations of electron microscopy images from mouse white matter [17]. These experiments do express internal gradients in the intra-axonal compartment. However, the 2D representation corresponds to infinitely long straight axons and thereby neglects all effects from the histology-documented longitudinal variations (diameter variation and tortuosity). Hence, any gradients that would be induced in the third dimension along the axons are not expressed.

In this work, we explore susceptibility-induced orientation-dependency of the SE and PGSE signals of myelinated axons at 7 T. We perform in silico experiments by integrating susceptibility-induced internal gradients in Monte Carlo diffusion simulations. With in silico experiments in individual axons, we study how different 3D morphological characteristics affect the orientation-dependency of the intra-axonal compartment. The characteristics have been quantified from XNH imaging of the corpus callosum (CC) of a vervet monkey brain [14], and include non-circular cross-sections, longitudinal diameter variations, and trajectory variations. With in silico experiments in densely packed axons, we study differences in orientation-dependency across the intra-axonal and extra-axonal compartments at different orientations. To validate our findings in real scans, we scan a cube of an ex vivo vervet monkey brain, and analyse the PGSE signals of the CC and the two cingula (CING-L and CING-R).

## 2 Results

### 2.1 Susceptibility-induced orientation-dependence of intra-axonal PGSE signal can distinguish morphological characteristics

We integrated susceptibility-induced internal gradients into Monte Carlo diffusion simulations in the intra-axonal compartments of realistic axon morphologies. The morphologies were quantified from an XNH volume of the CC of a vervet monkey brain. Our analysis reveals that different morphological characteristics cause different trends of susceptibility-induced orientation-dependency (axon w.r.t. **B_0_**) for the PGSE signal. We analyze both *S*(*b_des_* = 0) s/mm^2^ (corresponding to an SE signal and thereby a T2 effect), and diffusion tensor (DT) metrics fitted to b-values in different ranges. While the DT does not fully characterise the non-Gaussian diffusion in white matter, it does capture the principal anisotropy of the dMRI signal which is of our interest here. The morphological characteristics (longitudinal diameter variation, longitudinal trajectory variation, and cross-sectional eccentricity) are studied in five different configurations (C1-5) of increasing complexity; ranging from straight cylinders (Fig. 1a, C1), to real axons segmented from an XNH image (Fig. 1a, C5). First, we explore how the internal gradients induced by the different morphological configurations affect the signal at *b* = 0 s/mm^2^. This is done by normalizing the effective signal *S_eff_* (*b* = 0) (i.e. affected by internal gradients), w.r.t. the desired signal *S_des_*(*b* = 0) (i.e. unaffected by internal gradients and corresponding to the number of particles). Fig. 1b shows that each morphological configuration induces a unique trend of orientation-dependency. C1 expresses no orientation-dependency; in correspondence with the analytic model for straight coaxial cylinders [18]. C2 and C3 express the largest effect at angle(**B_0_**, **ẑ**) = 0 deg and angle(**B_0_**, **ẑ**) = 45 deg respectively. C4 expresses a combination of the effects observed for C2 and C3; in correspondence with the characteristics of this configuration in fact being a combination of the two. C5 expresses the strongest effect at angle(**B_0_**, **ẑ**) = 90 deg, and is affected more than any other configurations at any orientation.

**Figure 1:**
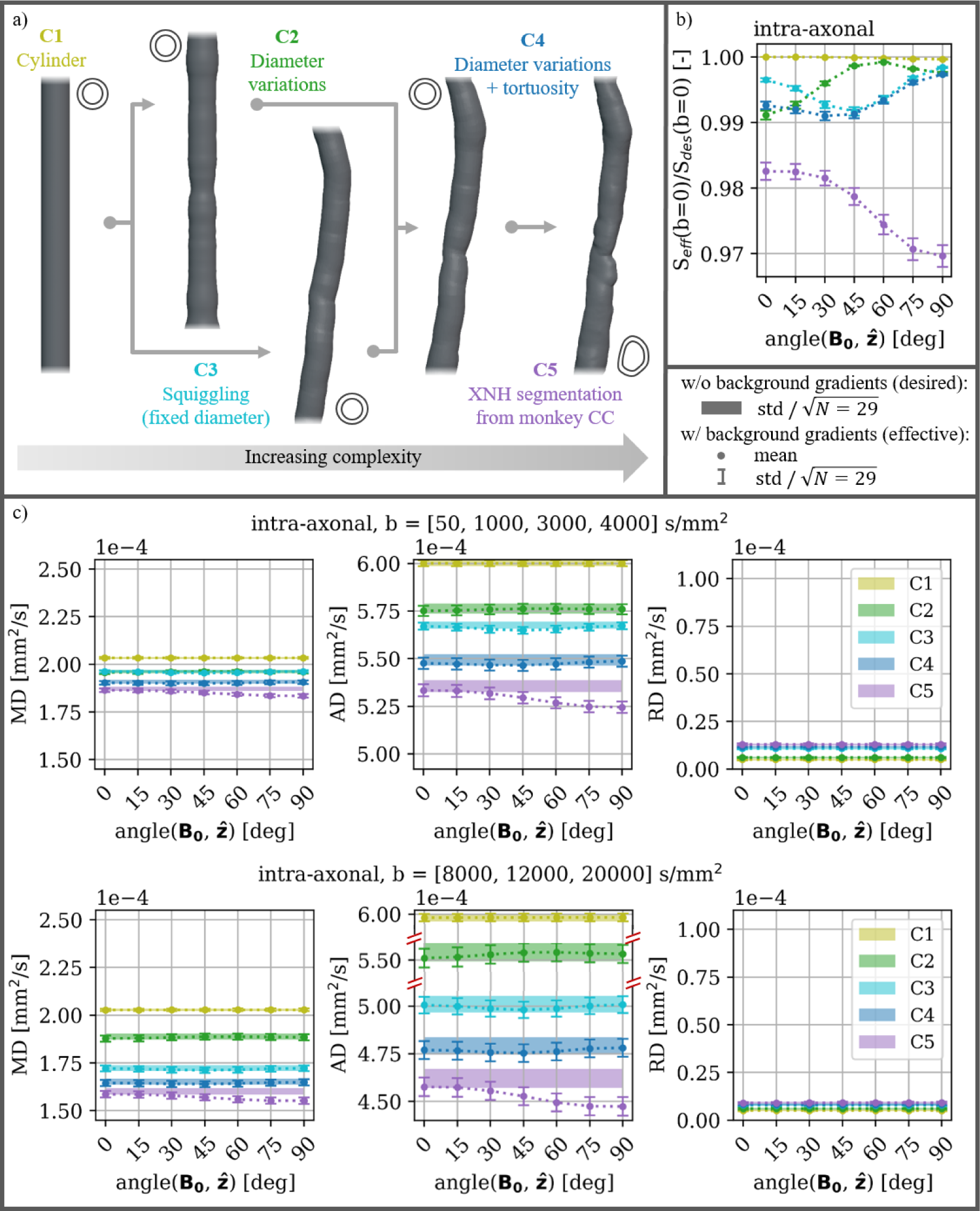
a) Schematic overview of the XNH-informed axon configurations. Axon lengths are cropped to 40 *µ*m for visualization purposes. The morphological characteristics (mean diameter, longitudinal diameter variation, and trajectory) were quantified from 29 axons segmented from an XNH-volume, and expressed in the five configurations (C1-5). C5: segmentations of real axons, C1: straight with the mean diameter of the corresponding C5-axons, C2: straight with longitudinal diameter variations of the corresponding C5-axons, C3: mean diameter and tortuous trajectories of the corresponding C5-axons, and C4: longitudinal diameter variations and tortuous trajectories of the corresponding C5-axons. The overall orientation of each axon is aligned with **ẑ**. **b)** The effect on the effective signal *S_eff_* measured relative to the desired signal *S_des_* at *b* = 0 s/mm^2^ as a function of orientation w.r.t. **B_0_**. This corresponds to an SE signal and thereby a T2 effect. **c)** Mean diffusivity (MD), axial diffusivity (AD), and radial diffusivity (RD) obtained from DT fits to lower b-values (upper row) and higher b-values (lower row - note the discontinuous y-axis on the AD plot). Markers and error bars indicate mean and deviation of the metric for *S_eff_* over experiments in the 29 axons. Bands indicate the deviation of the metric centered around the mean for *S_des_*. All y-axes have the same scale to emphasize the difference in the degree of orientation-dependency between the different metrics.

When fitting the DT to the PGSE signal, the same trends of orientation-dependency are observed for the mean diffusivity (MD) and the axial diffusivity (AD), while the radial diffusivity (RD) has no orientation-dependency (Fig. 1b vs. 1c). The orientation-dependencies are strongest for AD, and weaker for MD which is a combination of AD and RD. The difference in the effect on AD and RD shows that the susceptibility-induced internal gradients are causing an anisotropic effect on the PGSE signal, and that this anisotropy is orientation-dependent. By comparing DT metrics fitted at lower b-values ([50, 1000, 3000, 4000] s/mm^2^) with those fitted at higher b-values ([8000, 12000, 20000] s/mm^2^), an increased orientation-dependency is seen at higher b-values (Fig. 1c).

The different axonal configurations have unique effects on the DT metrics fitted to *S_des_*(*b*) for both b-value regimes. AD decreases with increasing configuration number; in correspondence with increasing obstructions in the form of longitudinal morphological variation in the axial direction. RD increases with increasing configuration number; in correspondence with added axonal tortuosity in the radial direction. MD is dominated by AD due to the elongated axonal morphologies (*λ*_1_ ≫ *λ*_2_*, λ*_3_ of the DT fits), and follows a similar decrease. Comparing DT metrics fitted at lower b-values with those fitted at higher b-values, shows a large decrease in the AD estimates for the morphological configurations expressing tortuosity. Meanwhile, C2 expresses only a small decrease, and C1 expresses no decrease. DT metrics for all morphological configurations show larger standard deviations at higher b-values. These results are in correspondence with previous work [14].

### 2.2 Susceptibility-induced orientation-dependency varies across tissue compartments and with the degree of axonal undulations

We now explore the effect of increasing the degree of undulation, and how the intra-axonal compartment is affected by internal gradients relative to the extra-axonal compartment. Undulation describes bending in a regular and repeating pattern, whereas tortuosity describes irregular bending. For this purpose, we construct numerical substrates of densely packed myelinated axons, where myelinated axons are constructed as coaxial helices on a hexagonal grid. The substrates resemble the C3 axons from the XNH-informed axon configurations (Fig. 1a), i.e. constant diameter and tortuous axon trajectories.

As the undulation amplitude increases, the direction of **e_1_** starts deviating from the axis of the helix (**ẑ**) (Sup. Fig. 9). Before including susceptibility-induced internal gradients in the signal, only the signal of the extra-axonal compartment has severe (>5 deg) deviation between **e_1_** and **ẑ**. When internal gradients are integrated in the signal, we see orientation-dependent deviation between **e_1_** and **ẑ** when *r_helix_* ≥ 4*µ*m. These deviations cause AD (the highest diffusivity of the DT) to not always be measured along the axial direction of these axons.

Without including internal gradients in the signal (*S_des_*), the different degrees of undulation (*r_helix_*) have different effects on the different DT metrics. AD decreases with *r_helix_*, and RD increases with *r_helix_*; in correspondence with increasing isotropy of the substrate. MD is dominated by AD when *λ*_1_ >> *λ*_2_*, λ*_3_, and follows a similar decrease with *r_helix_*.

The T2 effect caused by the susceptibility-induced internal gradients is analyzed by normalizing the effective signal *S_eff_* (*b* = 0) (i.e. affected by internal gradients), w.r.t. the desired signal *S_des_*(*b* = 0) (i.e. unaffected by internal gradients). Fig. 3c shows that the signals of the different tissue compartments have unique trends of dependency for both orientation and amplitude of the undulation. The substrate with *r_helix_* = 0.0 *µ*m expresses no intra-axonal effect at any orientation, no extra-axonal effect at angle(**B_0_**, **ẑ**) = 0 deg, and the largest extra-axonal effect at angle(**B_0_**, **ẑ**) = 90 deg; in correspondence with our expectations for the analytic model for straight coaxial cylinders [18]. An intra-axonal effect arises with increasing *r_helix_*. The extra-axonal compartment express minimal dependence on *r_helix_* at angle(**B_0_**, **ẑ**) = 0 deg, increasing dependence at angle(**B_0_**, **ẑ**) = 45 deg, and decreasing dependence at angle(**B_0_**, **ẑ**) = 90 deg. The signal from the combined compartments shows a combination of the two individual compartments.

**Figure 2:**
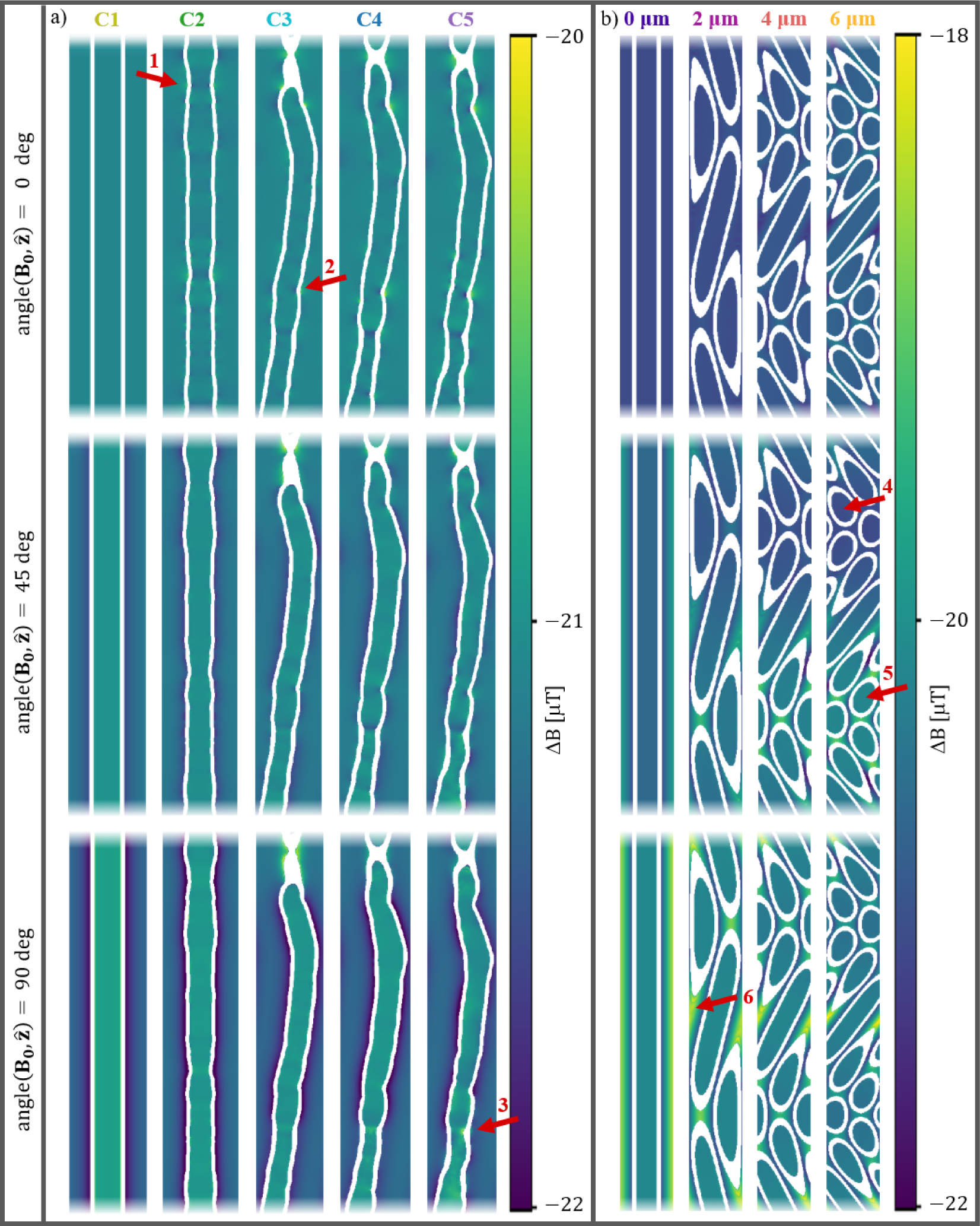
Cross-sections of the susceptibility-induced B-field changes at three different orientations between **B_0_** and the dominating orientation of the axons **ẑ**, i.e. angle(**B_0_**, **ẑ**) = [0, 45, 90] deg. To emphasize the field inhomogeneity in the compartments of interest (intra- and extra-axonal) the myelin compartment is hidden (shown in white), and the colour maps are cropped. **a)** C1-5 axons (see Fig. 1.a). Cropped to 50 *µ*m for visualization purposes. **b)** Helical axons (see Fig. 3.a) yz-cross-sections showing an entire period *λ* = 50 *µ*m of the periodic substrate. The arrows point to different examples of internal gradients. 1: caused only by diameter variation and most prominent at angle(**B_0_**, **ẑ**) = 0 deg. 2: caused only by tortuosity. 3: caused by the real axon morphology of the C5 and expressing a more complex pattern than what is observed at arrows 1 and 2. 4 and 5: attention to the difference in internal gradients induced along the helical axon for segments oriented differently w.r.t. **B_0_**. 6: more prominent gradients in the extra-axonal compartment compared with the intra-axonal compartment.

**Figure 3:**
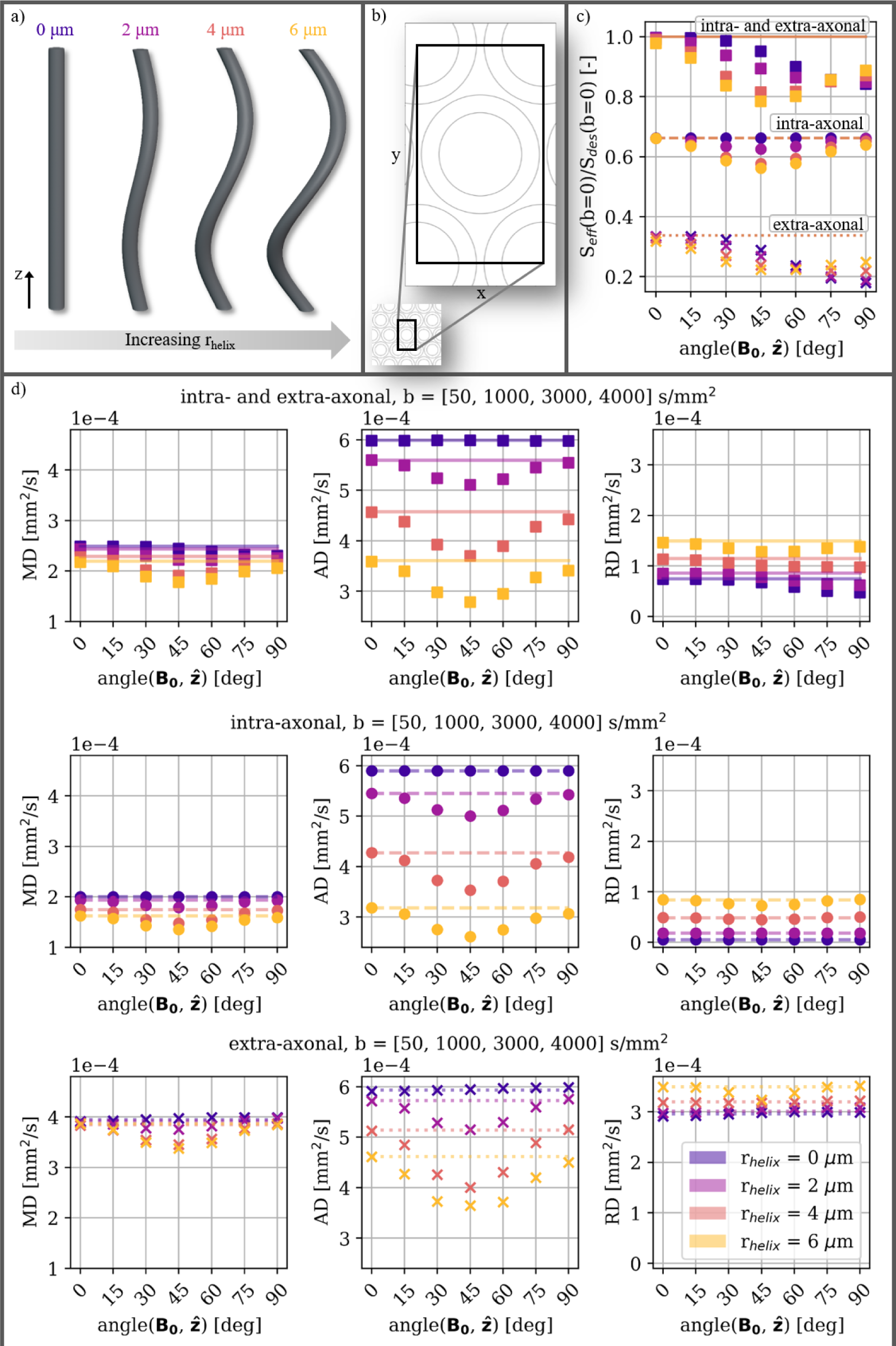
a) Helical axons with radii *raxon*=1.90 *µ*m, helical wavelength *λ*=50 *µ*m, and varying helical radii *r_helix_*. **b)** Cross-section of the unit voxel which is used to represent the hexagonally packed substrates. **c)** The effect on the effective signal *S_eff_* measured relative to the desired signal *S_des_* at *b* = 0 s/mm^2^ as a function of orientation w.r.t. **B_0_**. This corresponds to an SE signal and thereby a T2 effect. Different markers indicate the DT metrics fitted to *S_eff_* (*b*) for the different compartments. Lines indicate the DT metrics fitted to *S_des_*(*b*). **d)** Mean diffusivity (MD), axial diffusivity (AD), and radial diffusivity (RD) obtained from the DT model. Results for the combined intra- and extra-axonal compartments are shown in the upper row, for the intra-axonal compartment in the middle row, and for the extra-axonal compartment in the lower row. Markers indicate the DT metric fitted to *S_eff_* (*b*). Lines indicate the DT metric fitted to *S_des_*(*b*).

The DT metrics fitted to the individual compartments (Fig. 3.d) have the strongest orientation-dependency for AD, while it is weaker for MD. RD demonstrates orientation-dependency only when *r_helix_* = 6.0 *µ*m (the largest *r_helix_* tested here). This is also the substrate with the largest deviation between **e_1_** and **ẑ** (Sup. Fig. 9).

The DT metrics fitted to the combined signal of the intra- and extra-axonal compartments have noticeable orientation-dependent tissue-compartmental filtering due to the T2 effect at *b* = 0 s/mm^2^. This is based on the signals from the individual compartments showing different degrees of signal loss in Fig. 3.c. MD and RD have orientation-dependencies that are different from those observed for the individual compartments (minimum value around 45 deg), while more similar to the T2 effects (described above).

### 2.3 Susceptibility-induced orientation-dependence in ex vivo tissue

To test the orientation-dependency observed in silico, we performed ex vivo MRI scans of a tissue-block dissected from a perfusion-fixated vervet monkey brain, and scanned it at different orientations w.r.t. **B_0_**. Two acquisitions were made; for b-values up to 4000 s/mm^2^ in scan-series 1 (Fig. 12), and up to 20000 s/mm^2^ in scan-series 2 (Fig. 10, 11, and 4). To assess the impact in different tissue compartments, we utilize that higher b-values attenuate the signal originating from faster diffusing nuclei and thereby the extra-axonal compartment. Thus, we fitted a DT model to different ranges of b-values. As validated in Sup. Fig. 8, the signal from the highest range (*b* = [8000, 12000, 20000] s/mm^2^) originates from an isolated intra-axonal compartment. Meanwhile, the lower b-value ranges also reflect different degrees of the extra-axonal compartment. Orientation-dependency was quantified by ranking the best fitting trend (A-D) (see Sec. 4.4) based on the rescaled Akaike information criterion (ΔAIC). Overall, we found that the degree of orientation-dependence (quantified in the fitting parameter *p*_1_) and the magnitude (quantified in the fitting parameter *p*_0_) of the DT metrics decrease with increasing range of b-values. Meanwhile, the degree of relative orientation-dependency (*p*_1_*/p*_0_ 100) increases with the range of b-values (listed below). The degree of orientation-dependency in all ROIs is consistently largest for AD, and larger for CC than for CING. The difference in orientation-dependency of AD and RD indicates an anisotropic effect on the PGSE signal.

**Figure 4:**
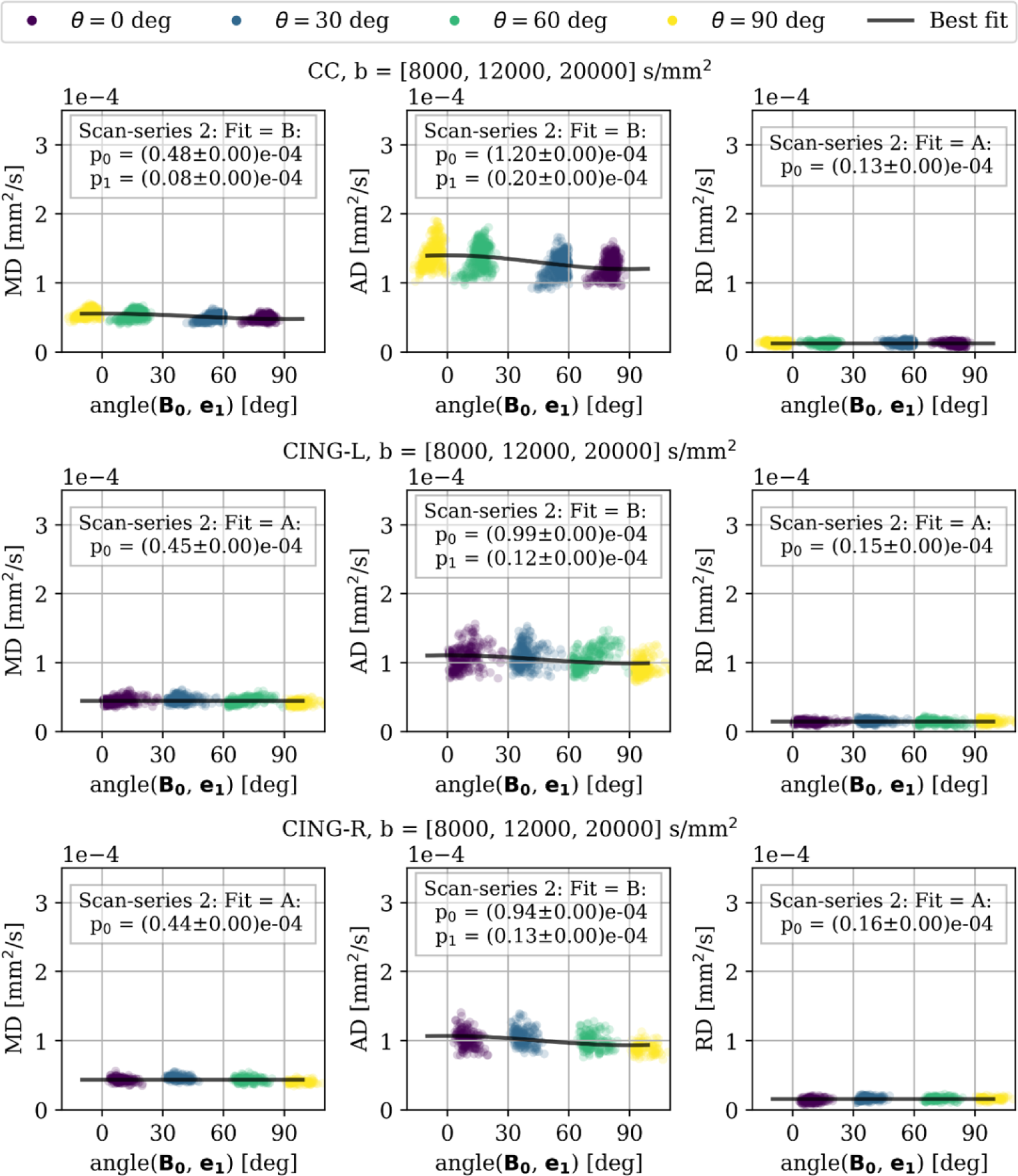
Scan-series 2. DT metrics fitted at high b-values ([8000, 12000, 20000] s/mm^2^) for CC (upper row), CING-L (middle row), and CING-R (lower row) of an ex vivo monkey brain scanned at 7 T. Markers are coloured according to which orientation *θ* the scan was acquired at, and plotted along with the best fit based on lowest AIC-value. Orientation-dependency is observed for AD of all ROIs, and for MD of CC. The degree of orientation-dependency is stronger for CC than for CING-L and CING-R. The difference in effect on AD vs. RD shows that the PGSE signal is affected anisotropically. ΔAIC and ΔRMSE for all DT metrics and all fitted models are listed in Sup. Tab. 3

At high b-values ([8000, 12000, 20000] s/mm^2^, scan-series 2 in Fig. 4), where the signal is dominated by the intra-axonal compartment (validated in Sup. Fig. 8), CC demonstrates orientation-dependency of trend B for both MD and AD, and no orientation-dependency (trend A) for RD. This agrees with our in silico findings for the C5 axons which are also quantified from monkey CC. Meanwhile, CING-L and CING-R demonstrate orientation-dependency of trend B only for AD, and no orientation-dependency (trend A) for MD or RD. The estimated degrees of relative orientation-dependency are 16% for MD and 17% for AD in CC, and 12-14% for AD in CING.

At intermediate b-values ([4000, 8000, 12000] s/mm^2^, scan-series 2 in Sup. Fig. 11), CC, CING-L, and CING-R demonstrates orientation-dependency of trend B for both MD and AD, and no orientation-dependency (trend A) for RD. However, both trend A and B show substantial support for MD in CING-R with ΔAIC<2, and also trend B show some support for MD in CING-L with 4 ≤Δ*AIC*≤7. The estimated degrees of relative orientation-dependency are 15% for MD and 13% for AD in CC, and 12-12% for MD and 9-10% for AD in CING.

At low b-values ([50, 1000, 3000, 4000] s/mm^2^, scan-series 2 in Sup. Fig. 10 and scan-series 1 in Sup. Fig. 12), all ROIs demonstrate orientation-dependency of trend B for both MD and AD, and no orientation-dependency (trend A) for RD. In scan-series 2, the estimated degrees of relative orientation-dependency are 7% for MD and 8% for AD in CC, and 5-5% for MD and 5-6% for AD in CING.

## 3 Discussion

With in silico experiments, we show that the susceptibility-induced orientation-dependency trends for the intra-axonal SE and PGSE signals are specific to the morphological characteristics expressed in an axon population. Both the impacts of cross-sectional and longitudinal morphological variations prove crucial. Compared to previous 2D modelling [17], our 3D modelling thereby provides additional insight into susceptibility-induced orientation-dependency of myelinated axons. Interestingly, our findings show potential for a novel sensitivity to cross-sectional eccentricity of myelinated axons. This sensitivity is not attainable with the theoretical dMRI signal (*S_des_*), due to the limited spatial resolution averaging over an ensemble of axons, and thereby smearing out the eccentricity of the individual axons. However, we demonstrated that the sensitivity to eccentricity can be obtained by leveraging the susceptibility-induced orientation-dependency. Furthermore, we show that the intra-and extra-axonal compartments can be affected differently by these internal gradients, and thereby cause orientation-dependent filtering of the signals from those compartments.

With ex vivo experiments in monkey (CC, CING-L and CING-R), we reproduced the orientation-dependency trend observed only for the in silico axon population expressing the full range of morphological characteristics (longitudinal diameter variations, tortuosity, and cross-sectional eccentricity - i.e. real axons segmented from XNH volumes). Hence, our results contradict the more simple representations of myelinated axons without longitudinal morphological variation.

We show that AD demonstrates orientation-dependency while RD remains stable. We attribute this anisotropic effect of the PGSE signal to arise from the cross-terms between the external and internal gradients. Previously, orientation-dependency of dMRI has only been documented across different white matter regions where differences in microstructure across regions cannot be ruled out as a contributing factor [19][20][21].

The orientation-dependency observed in this study is further reason to take caution when comparing signals from different regions of myelinated axons. This is relevant both for axon populations oriented differently w.r.t. **B_0_**, and for axon populations expressing different morphological characteristics. If effects of internal gradients are not incorporated in the applied modelling, the certainty of the model estimates may be limited due to the orientation-dependency. This becomes only more relevant with higher **B_0_** strengths.

### 3.1 Linking axon morphology characteristics to orientation-dependency

With in silico experiments, we show that different morphological characteristics of myelinated axons (longitudinal diameter variations, tortuous trajectories, and cross-sectional eccentricity) introduce different susceptibility-induced orientation-dependency trends of the intra-axonal SE and PGSE signals. The different morphological configurations (C1-5) of increasing complexity are seen in Fig. 1.a. In this section, we focus on the results for SE experiments shown in Fig. 1.b where the trends are observed most clearly.

C1 not expressing any orientation-dependency of intra-axonal signals is explained in the analytic solution for the coaxial cylinder [11][12][13], where no gradients are induced in the intra-axonal compartment.

C2 expressing the largest effect at angle(**B_0_**, **ẑ**) = 0 deg may be explained by Eq. 4, where the morphological variations in the direction of **B_0_** have the greatest influence on the resulting B-field. Because C2 has circular cross-sections, the morphology with the greatest influence when approaching angle(**B_0_**, **ẑ**) = 90 deg will resemble the coaxial cylinder (i.e. C1). Meanwhile, the morphology with the greatest influence when approaching angle(**B_0_**, **ẑ**) = 0 deg is the longitudinal diameter variations.

C3 and the helixes having the largest effect at angle(**B_0_**, **ẑ**) = 45 deg may be explained by the distribution of angles between **B_0_** and discretised longitudinal axon segments (angle(**B_0_**, axon segments)) (see Sup. Fig. 7). When angle(**B_0_**, **ẑ**) = 0 deg, the distribution of angle(**B_0_**, axon segments) is relatively narrow due to the axons’ overall alignment with **ẑ**. When angle(**B_0_**, **ẑ**) = 90 deg, the distribution of angle(**B_0_**, axon segments) is again relatively narrow due to the axial symmetry around 90 deg. When angle(**B_0_**, **ẑ**) = 45 deg, angle(**B_0_**, axon segments) has the widest distribution extending all way from angle(**B_0_**, **ẑ**) = 0 deg to 90 deg. This orientation thereby captures the largest variety of induced field inhomogeneity.

C4 demonstrating a combination of the effects of C2 and C3, may be explained by the morphological characteristics expressed in C4 being the combination of those expressed in C2 and C3.

Interestingly, the trend of C5 stands out significantly from that of C4. The axons of C5 are the actual segmentations of real axons, and the only difference from C4 is eccentric cross-sections. C5 stands out by demonstrating the largest effect at angle(**B_0_**, **ẑ**) = 90 deg, a larger degree of orientation-dependency overall, and a significant negative offset of the signal compared with C1-4. These effects may be explained by the eccentricity which due to the associated irregularities induces increased field inhomogeneity at all orientations, and especially at angle(**B_0_**, axon segments) = 90 deg. For C1-4 the morphology with the greatest influence when approaching angle(**B_0_**, **ẑ**) = 90 deg will resemble the coaxial cylinder due to their circular cross-sections. Furthermore, the spatial variations in susceptibility associated with the eccentric cross-sections occur on a short length scale compared to the variations due to diameter variations and tortuosity, and the associated phase accumulations are therefore less likely to be cancelled out.

When acquired at a single orientation w.r.t. **B_0_**, the dMRI signal is not sensitive to eccentricity due to the limited spatial resolution averaging over an ensemble of axons. However, by acquiring scans at multiple orientations, the orientation-dependency trend appears to be sensitive to eccentricity.

### 3.2 Orientation-dependent filtering of tissue compartments

We show that susceptibility-induced internal gradients can give rise to compartment-specific effects, and thereby orientation-dependent filtering (or weighting) of the tissue compartments. The simplest example of this is seen for the substrate with *r_helix_* = 0 *µ*m (i.e. straight coaxial cylinders) in Fig. 3. Although neither the intra-axonal nor the extra-axonal compartments demonstrate orientation-dependency of the RD, the combined signal does (Fig. 3.d, rightmost column). This can be explained by orientation-dependent filtering of the signals at *b* = 0 s/mm^2^ (Fig. 3.c). While the signal of the intra-axonal compartment does not appear affected by internal gradients at angle(**B_0_**, **ẑ**) = 90 deg, the extra-axonal compartment does. Hence, at this orientation, the combined signal will have a smaller contribution from the extra-axonal compartment compared with the orientation angle(**B_0_**, **ẑ**) = 0 deg. Since the extra-axonal RD is higher than the intra-axonal RD, this results in a lower combined RD. This generalizes to the substrates with *r_helix_ >* 0 *µ*m. We expect this concept to generalise to intra- and extra-axonal compartments with other morphological configurations.

### 3.3 With great b-value comes great filtering

We show that AD of the intra-axonal signal decreases through the morphological configurations C1-5; in correspondence with increasing obstructions in the axial direction caused by the morphological characteristics (Fig. 1.c). The dependence on obstructions in the axial direction increases significantly at higher b-values. Especially the configurations expressing tortuosity (i.e. C3-5) are affected. For *S_des_*(*b*) this is solely due to the higher b-values filtering out the faster diffusing components (i.e. sections of the axons parallel with the external gradient). In continuation, this leads to filtering of which morphological aspects are represented in the signals. Morphological aspects leading to lower diffusivity will thereby dominate the signal. Because such aspects are likely to be correlated with characteristics of internal gradients, we expect *S_eff_* (*b*) to likewise reflect a subset of internal gradients and thereby influence the filtering. Furthermore, the stronger external gradients at the higher b-values will lead to a stronger influence by the cross-terms between the internal and external gradients. Since the internal gradients are anisotropic, so are the cross-terms. This may lead to additional directional filtering of the signal (discussed further in Sec. 3.4).

Our ex vivo experiments reproduced the orientation-dependency trend observed only for the in silico axon population expressing the full range of morphological characteristics (longitudinal diameter variations, tortuosity, and cross-sectional eccentricity). We observe an increase in orientation-dependency at higher b-values where the signal is dominated by contributions of lower diffusivity, and therefore assumed to be dominated by the intraaxonal compartment. This may indicate that the subset of internal gradients associated with the contributions of lower diffusivity causes more pronounced susceptibility-induced effects. This is likely a consequence of a higher degree of non-linearity of these internal gradients. However, disentangling the combined effects of internal and external gradients is a complex matter.

### 3.4 Anisotropy of cross-term contributions

With both in silico and ex vivo experiments, we show that the PGSE signal in white matter is affected anisotropically by susceptibility-induced internal gradients. How the internal gradients affect the PGSE signals depends on the anisotropic cross-terms between internal and external gradients, and the anisotropic apparent diffusivity [1][3]. The DT assumes the signal to follow exp(*b_des_D*) [22][23], when in fact the signal follows exp(*b_eff_ D*). This means that any diffusion weighting deviating from *b_des_* will be reflected in the fitted variable *D*. Hence, any additional diffusion weighting from internal contributions will lead to an underestimation of *D* in the given direction - and vice versa. If we consider the hexagonally packed axons with *r_helix_* = 0 *µ*m (Fig. 3), we see that whenever the axons are not parallel with the applied field, internal gradients will be induced in the extra-axonal space. This is in correspondence with analytic solutions for the straight coaxial cylinder [11][12][13]. The gradients will be induced perpendicular to the axis of the axons, meaning that any line parallel with the axon will be an isoline. As expected, the AD (in this case modelling diffusion along the isolines of the internal gradients) shows no impact from internal gradients. When *r_helix_ >* 0 *µ*m the isolines of the internal gradients are no longer straight and parallel with the AD. In this case, the degree of orientation-dependence increases with *r_helix_*. The RD, however, is modelling diffusion perpendicular to the isolines of the internal gradients for *r_helix_* = 0 *µ*m, but still shows no impact by internal gradients. We believe this is due to the much lower diffusivity in the plane perpendicular to the axons, as this allows much less probing of the internal gradients.

### 3.5 Expected impact of susceptibility-induced biases in dMRI modelling

We studied the orientation-dependency of signals from myelinated axons. Naturally, demyelination [24][25][26] would affect the results. While demyelination is best detected with other MRI modalities such as T2 and FLAIR [27][28], and the primary effect is assumed to be on the extra-axonal diffusion processes and increased exchange across compartments, our findings suggest that also intra-axonal DT metrics can be affected directly through the change in internal gradients.

We show that axon diameter variations can cause significant intra-axonal signal loss when the axons are oriented parallel with **B_0_** (C2 in Fig. 1). Axonal beading (and thereby amplified diameter variations) has been observed in various diseases such as multiple sclerosis [24], ischemic stroke [29], and traumatic brain injury [30][31]. We expect such pathomorphology to influence the orientation-dependency trends of *S_eff_* (*b* = 0) and the AD.

We document morphology- and orientation-dependency of AD. Several models for PGSE signals are based on a fixed value for the axial diffusivity. This may lead to a bias across different brain regions because the different regions express different axon morphologies and orientations. By assuming a fixed value for the axial diffusivity, the susceptibility-induced variation of axial diffusivity may therefore be reflected in the additional model parameters instead. Examples are axon diameter estimations with ActiveAx [32] and with SMT [33], and orientation dispersion and density estimates with NODDI [34]. However, some tolerance to imprecise fixation of axial diffusivity has been documented for the SMT model [35].

### 3.6 Limitations

#### 3.6.1 Field strength and sequence-parameters

The degree of susceptibility-induced internal gradients increases with increasing field strength. Here, we studied the effects at 7 T. We expect more severe effects at higher field strengths. With a tendency of moving towards higher field strength due to the increased SNR, understanding the effects of internal gradients become only more relevant.

We show the results for one set of PGSE sequence-parameters. However, the results for both *S_des_*(*b*) [36][37][14][38] and *S_eff_* (*b*) [39] will depend on the choice of sequence-parameters.

#### 3.6.2 In silico experiments

Due to the high memory usage when computing the susceptibility-induced internal gradients at adequate spatial resolution, our in silico experiments were limited to only very small unit voxels. This did not enable representation of a realistic packing of the axons. Novikov et al. [39] showed that, compared to random packing, square packing limits the impact of internal gradients in 2D simulations due to the repetitive pattern of the induced internal gradients. We expect a similar effect for the hexagonal packing investigated here. A more realistic packing could be obtained by assuming the interaction between the fields of individual axons to be negligible compared to the interaction between the individual axon and **B_0_**. This could be implemented by computing the internal gradients of individual axons, and then adding these contributions to an arbitrary grid while allowing super-position only in the extra-axonal compartment.

Myelin is believed to be the main contributor to internal gradients in white matter [8]. We studied the effect of different axon and myelin morphology but did not consider effects from nodes of Ranvier. Such variations in myelin are expected to induce further inhomogeneity. We neglected dipolar effects arising from the anisotropically organized lipids of myelin [13], which could contribute to a magic angle effect [40]. We approximated the susceptibility values for myelin, intra-axonal and extra-axonal compartments based on those of fat and water. Deviation in the differences of susceptibility across compartments would influence the results. Larger differences would cause stronger field inhomogeneity and vice versa.

Due to the differences in the internal gradients induced in the individual tissue compartments, we expect that exchange across the myelin has an impact on the effective signal. We did not model such effects here.

#### 3.6.3 Ex vivo experiments and relation to in vivo systems

Several aspects make it infeasible to directly transfer our ex vivo findings to in vivo predictions. The complete impact of tissue fixation is unknown, but both microstructural and diffusion properties undergo changes compared to when the tissue is in its in vivo state. The tissue scanned here, was perfusion fixated with formaldehyde which is the most well-established chemical fixation agent for MRI [41]. However, chemical fixation with aldehydes may in some cases compromise the myelin structure by loosening and splitting of adjacent membrane layers [42]. This would change the bulk susceptibility properties of the myelin, and hence, the susceptibility-induced internal gradients. Although fractional anisotropy is reported as stable under these conditions, D’Arceuil et al. [43] report up to 80% decrease in the apparent diffusion coefficient in white matter due to formalin fixation. Birkl et al. [44] report a 4% decrease in water content, and a >25% decrease in T2 values in white matter due to formalin fixation. By rinsing the tissue with saline prior to scanning (as done here), T2 values can be partly restored [45]. Based on these deviations between in vivo and ex vivo tissue, we expect stronger susceptibility effects in vivo due to 1) higher diffusivity, 2) longer TEs, and 3) more structured myelin.

## 4 Methods

### 4.1 Monkey brain tissue

One brain was prepared as part of the synchrotron experiment described in Andersson et al. [14] to image axons in nanometer image resolution. The second brain was used for the dMRI experiments. The tissue comes from two perfusion fixed female and age matched (32 month) vervet (Chlorocebus aethiops) monkey brains, obtained from the Montreal Monkey Brain Bank. The monkeys, cared for on the island of St. Kitts, had been treated in line with a protocol approved by The Caribbean Primate Center of St. Kitts. The brains had previously been stored and prepared according to Dyrby et al. [46].

### 4.2 In silico experiments

In silico DWI experiments were carried out for two branches of axon-mimicking numerical substrates: 1) isolated axons expressing different combinations of morphological characteristics quantified from axons previously segmented from synchrotron X-ray nano-holotomography (XNH) volumes [14], and 2) hexagonally packed helical axons (coaxial cylinders). Monte Carlo diffusion simulations were carried out with surface mesh representations of the numerical substrates. For each mesh representation we generated a three-compartment volume segmentation (intra-axonal, extra-axonal and myelin) as a voxelized representation from where we computed the susceptibility-induced internal gradients. The internal gradients were integrated into the Monte Carlo simulations by including the phase accumulation caused by the internal gradients when computing the DWI signal.

#### 4.2.1 XNH-informed axon configurations (intra-axonal compartment)

To study how different morphological characteristics of axons affect the susceptibility-induced internal gradients and the related SE and PGSE signals, we conducted simulations for 29 segmentations of real axons, and four different configurations of morphological characteristics quantified from each of these (Fig. 1a). The quantified characteristics are mean axon diameter, longitudinal diameter variation, and axonal trajectory. Expressed in the five configurations (C1-5) we have C5: segmentations of real axons, C1: mean diameter with straight axon trajectories (cylinders), C2: longitudinal diameter variations with straight axon trajectories, C3: mean diameter and tortuous axon trajectories, and C4: longitudinal diameter variations and tortuous axon trajectories. All axons are aligned with the z-axis.

The axons were segmented from a 3D XNH volume of the splenium of a 32-month old vervet monkey brain. The volume was acquired with 75 nm isotropic voxel size and 153.6 µm cylindrical field-of-view at beamline ID16A of the European Synchrotron. Fifty-four axons with a minimum length of 120 µm were semi-manually segmented in the native image resolution, after which their trajectories were determined, allowing for a quantification of longitudinal diameter variations. Local axonal diameters were quantified at 150 nm intervals and defined as the equivalent diameter – the diameter of a circle with the same area as the axonal cross-section in the plane perpendicular to the local trajectory, as in [16]. The XNH image acquisition, segmentation and analysis is fully described in [14]. To produce triangulated surface meshes for simulations, the binary segmentations were first meshed using the isosurface() function in Matlab R2020, and then processed with the Corrective Smooth Modifier in Blender (RRID:SCR_008606) for reduction of high-frequency changes and irregularities. The meshes were decimated in Blender to reduce triangle density, while preserving the mesh volume, to reduce the computational burden of the simulations.

For these simulations, the main direction of each axon was calculated using a principal-component analysis of its trajectory, after which it could be rotated into the z-axis. 29 axons were then selected based on an upper limit of the volume of their bounding boxes; in the interest of feasibility of computing the corresponding internal gradients (described in 4.2.3). These make up the C5 substrates. Axonal configurations C1-C4 were first modelled as consisting of z-aligned cylindrical segments, after which one or more of the following properties from the C5 axons was inherited by deforming the cylindrical segments in the appropriate position along the axonal trajectory: mean axon diameter, longitudinal diameter variation, tortuous trajectories. Importantly, although the axonal cross-sections in the plane perpendicular to the local trajectory could be non-circular and non-symmetrical for the C5 configuration, this property was not inherited by C1-C4 for which the axonal cross-sections were circular.

To compute the susceptibility-induced internal gradients, myelin compartments were constructed artificially by expanding the axon mesh. The expansion was performed by translating each vertex of the mesh a distance *width_myelin_* = *mean*(*r_axon_*)*/g*, where *mean*(*r_axon_*) is the mean radius of a given axon, and *g* = 0.7 is the g-ratio, in the direction away from the centreline of the original mesh. Each new mesh was then re-meshed, re-triangulated, and decimated in Blender (RRID:SCR_008606) to ensure the quality of the mesh-triangulation.

#### 4.2.2 Hexagonally packed helical axons (intra- and extra-axonal compartment)

To study how the susceptibility-induced internal gradients of tortuous axons affect the signal in the intra-axonal and extra-axonal compartments relative to each other, we generated numerical mesh substrates of densely packed axons (Fig. 5 and 3a). Packing and undulation was designed with periodic boundary conditions for optimal memory usage during computation of the internal gradients (described in 4.2.3).

**Figure 5:**
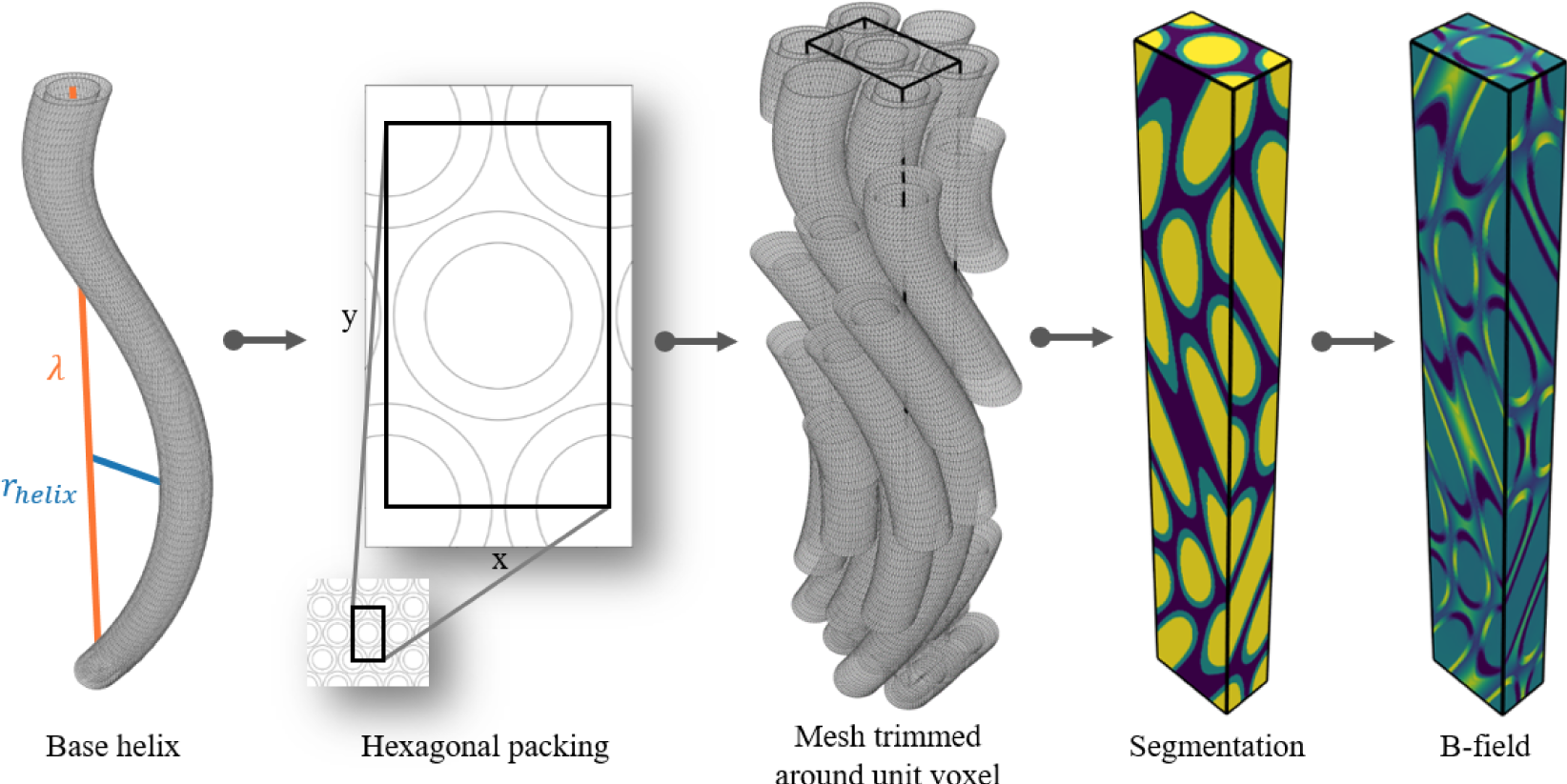
Pipeline for generating hexagonally packed helical axon substrates. A base axon is generated as a surface mesh, and defined by an axon radius *raxon*, a g-ratio, a helix wavelength *λ* (along **ẑ**), and a helix radius *r_helix_*. The hexagonal packing is defined by an extra-axonal volume fraction (EAVF), and *raxon*. Copies of the base axon are arranged according to the hexagonal packing, and the meshes are trimmed around the unit voxel of the hexagonal packing for computational efficiency. A compartment-wise segmentation is obtained as a voxel-representation of the surface mesh (intra-axonal in yellow, myelin in green, and extra-axonal in blue). B-fields are then computed by assigning magnetic susceptibilities to each compartment of the segmentation, and solving for **B_0_** applied in different directions angle(**B_0_**, **ẑ**).

The myelinated axons (coaxial cylinders) are organized with hexagonal packing in the xy-plane (cross-sectional to the axons), and helical undulation along the z-axis (i.e. along the primary axon axis). The hexagonal packing and helical undulation impose translation-symmetry in all dimensions and enables the substrate to be fully described from a volume containing a single period along each dimension. We define this volume as the unit-voxel. The side lengths (*l_x_, l_y_, l_z_*) of the unit-voxel are given by

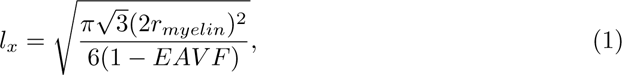

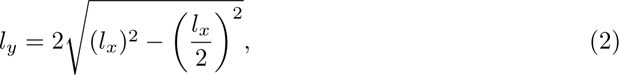

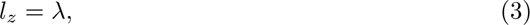

where *r_myelin_* is the outer radii of the myelinated axons in the xy-plane, *EAV F* is the extra-axonal volume fraction, and *λ* is the wavelength of the undulation along the z-axis. Substrates were generated with extra-axonal volume fraction *EAV F* = 0.2 in accordance with [47].

Myelinated axons were modelled as co-axial tubes with an outer radius *r_myelin_* = 1.9 *µ*m representing the radius of the myelin, an inner radius *r_axon_* = *r_myelin_ g* representing the radius of the axon, where *g* = 0.7 is the g-ratio (as in other studies [48][49]). Axonal trajectory undulation is modelled as a helical trajectory with wavelength *λ* and radius *r_helix_*. A surface mesh-representation of the axons was composed by stacking ring modules. The undulation causes axons to cross in and out of the unit-voxel. To save computation time during segmentation (described in 4.2.3) and Monte Carlo diffusion simulations (described in 4.2.4), the meshes are cropped module-wise around the unit-voxel.

#### 4.2.3 Computation of susceptibility-induced internal gradients

To compute the susceptibility-induced internal gradients of the numerical substrates, we computed how the 3D distribution of magnetic susceptibilities (based on the three tissue compartments) perturbs an externally applied magnetic field (B-field). This is infeasible to do analytically for all but simple and analytically described geometries. To solve B-fields for the complex geometries presented here, we therefore implemented a finite-difference (FD) method.

The FD method requires a voxel-representation of the substrate, i.e. the volume of the unit-voxel partitioned into voxels. This was obtained with a ray-casting-based [50] in-house GPU-compatible method. From the surface mesh-representation, the method segments the volume within the unit-voxel into three compartments (intra-axonal, extra-axonal, and myelin). All voxels in the myelin compartment of the voxel-representation are strictly contained within the myelin compartment of the surface mesh-representation; i.e. no myelin-voxels are transcending the meshes. This is crucial for preventing the B-field in the myelin compartment from contaminating the particles diffusing in the intra- and extra-axonal compartments. The resolution of the segmentations is a trade-off between descriptive accuracy of the morphology, and memory usage when computing the B-fields.We applied a desired resolution of *res_desired_* = 0.0655 *µ*m. This resolution had to be slightly modified (±0.0001 *µ*m) to match the unit-voxel of the individual substrate.

Exact susceptibility values for myelin, intra-axonal and extra-axonal compartments are unknown. We approximate the susceptibility of the intra- and extra-axonal compartments to that of water, i.e. *χ_water_* = *χ_intra_* = *χ_extra_* = 9.04 10*^−^*^6^ [51], and the susceptibility of the myelin compartment to that of fat, i.e. *χ_fat_* = 7.79 10*^−^*^6^ [51] with a myelin water fraction (MWF) of 0.15 in accordance with findings for MWFs for normal-appearing white matter (NAWM) in [52][53] by computing a weighted sum of the individual values, i.e. *χ_myelin_* = *MWF χ_water_* + (1 *MWF*) *χ_fat_* = 7.98 10*^−^*^6^.

The implementation of the FD method is inspired by Bhagwandien et al. [18][11], and applied to iteratively solve the magnetic scalar potential **Φ** from

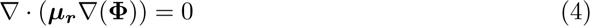

where ***µ_r_*** = ***χ*** + 1 is the volumemetic distribution of magnetic permeability of the substrate relative to that of vacuum. From **Φ** the magnetic field Δ**B** is then computed by

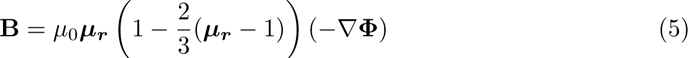

where *µ*_0_ is the magnetic permeability of vacuum. B-fields were computed for *B*_0_ = 7 T at different orientations angle(**B_0_**, **ẑ**) = (0, 15, 30, 45, 60, 75, 90) deg between the primary orientation axis of the axons **ẑ** and **B_0_**.

Due to the anisotropic organization of the lipids within the myelin, the B-field perturbations can be divided into an isotropic contribution and an anisotropic contribution [12], where the isotropic contribution comes from the bulk of the myelin, while the anisotropic contribution comes from the dipole effects of the lipids. The effect of the isotropic contribution is dominating when considering only the intra- and extra-axonal compartments and no permeability, and scales stronger with increasing field strength. The latter fulfils our requirements.

#### 4.2.4 Monte Carlo diffusion simulations and integration of internal gradients

To generate dMRI signals from the numerical substrates, we first performed Monte Carlo diffusion simulations to obtain the diffusion trajectories of particles restricted by the surface mesh-representations of the substrates. Then, we computed the resulting dMRI signals taking into account both the (diffusion encoding) external gradients and the (susceptibility-induced) internal gradients.

The diffusion trajectories were obtained for the surface mesh-representations of the numerical substrates with the MC/DC Simulator [54] with diffusion coefficient *D*_0_ = 0.6 *µ*m^2^*/*ms (as applied for ex vivo elsewhere [32]), step length *l_step_* = 0.11 *µ*m, and particle density *ρ_particles_* = 15 *µ*m*^−^*^3^ (corresponding to between 7360 and 23271 intra-axonal particles per XNH-informed substrate, and 14177 extra-axonal and 27786 intra-axonal per hexagonally packed substrate). Signal from the myelin compartment was neglected due to its short T2 (~20 ms) and T2* (10 ms) [55][56] compared to the applied TE. Particle initialisation was done only in the intra- and extra-axonal compartments, and permeability was set to zero.

For the XNH-informed axon configurations, particle initialisation was only done in the intra-axonal compartment. Particles were initialised at a minimum distance of 20 *µ*m from the closed ends to avoid probing these.

The simulation experiment used the same acquisition setup (b-values, b-vectors, TE) as in the ex vivo experiment (described in 4.3.2). The experiment was repeated at different orientations w.r.t. **B_0_** at angle(**B_0_**, **ẑ**) = (0, 15, 30, 45, 60, 75, 90) deg.

The dMRI signal was computed from the simulated diffusion trajectories by accumulating both the phase contribution from the external gradients of the PGSE sequence, and the phase contribution from the internal gradients caused at a given *θ_z_*. Given by

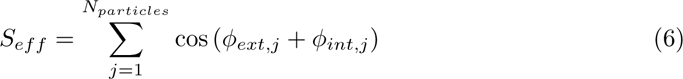

where *ϕ* is the temporally accumulated phase of a particle computed by

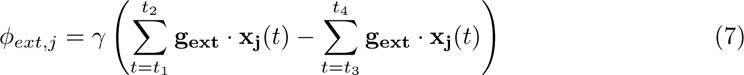

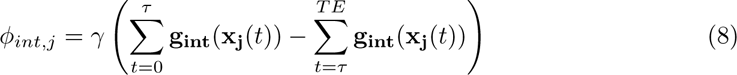

where **g_ext_**is the external gradient field, **g_int_** is the internal gradient field, *τ* = *TE/*2 is the inversion time, *t* is time, and *t*_1_, *t*_2_ = *t*_1_ + *δ*, *t*_3_ = *t*_1_ + Δ, and *t*_4_ = *t*_3_ + *δ* are the temporal boundaries of the external gradients, and *δ* is the duration of the external gradients.

### 4.3 Ex vivo MRI experiments

A tissue block of (~17 mm)^3^ was dissected from a whole perfusion fixated vervet monkey brain, and scanned at different orientations w.r.t. **B_0_** to validate the orientation-dependencies observed for the numerical substrates.

#### 4.3.1 Rotation device

The tissue block was placed in a rotation device as shown in Fig. 6. The rotation device was tailored for the RF-coil, such that the centre of the bowl is the centre of the coil when placed on the associated foundation. The sides of the bowl has the geometry of a hollow coaxial cylinder, which in the ideal case induces no internal gradients inside the bowl [11]. Orientation of the bowl was controlled with 30 deg increments according to the foot of the bowl. The tissue sample was moulded in 2% PBS-agar [57] to stabilize it mechanically in the bowl. The bowl was sealed off using Parafilm®and nitrile rubber to avoid dehydration during scanning. The bowl was placed on the foundation to control the rotation. The orientation of the brain w.r.t. **B_0_** is measured as the angle between **B_0_** and the longitudinal fissure.

**Figure 6:**
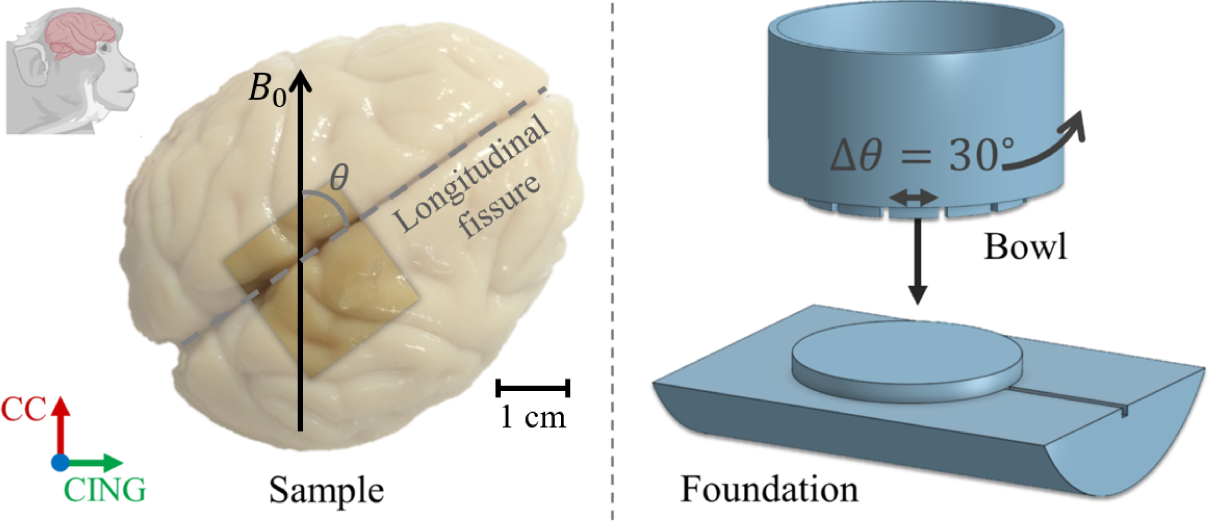
Ex vivo experimental set-up. **a)** Indication of the dissected tissue block, and how the orientation of the tissue w.r.t. **B_0_** was defined based on the longitudinal fissure. **b)** Rotation device tailored for the RF-coil. Some figure elements in a) were created with BioRender.com.

**Figure 7:**
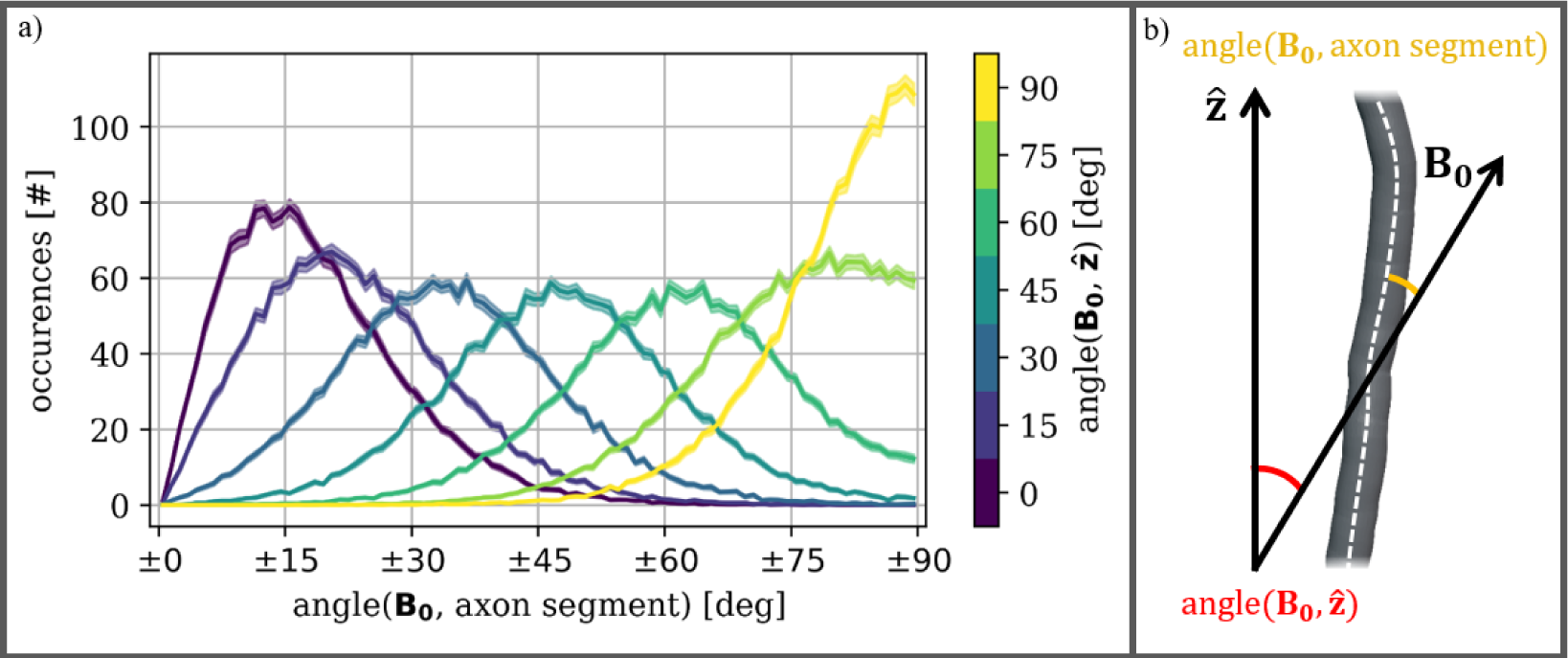
a) Distributions of angles between **B_0_** and longitudinal axon segments of C3, as they appear at different orientations between **B_0_** and the overall orie<u>ntation</u> of the axons **ẑ**. Lines are the mean distribution over all 29 C3 axons. Margins are means 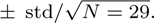. **b)** Illustration of how the angles in a) are measured for the C3 axons.

#### 4.3.2 MRI scan parameters

Two scan-series were acquired. Within each series, the scan-protocol (described below) was repeated for different orientations w.r.t. **B_0_** at *θ* = (0, 90, 30, 60) deg (Fig. 6). The order was pseudorandomized in order to exclude temporal bias in the analysis. Field-of-view was aligned with the longitudinal fissure at each scan.

Scans were performed on a 7 T Bruker Biospec 70/20 preclinical MRI scanner with maximum gradient strength of 600 mT/m, and a 1H transmit-receive volume quadrature RF-coil with inner diameter of 40 mm. Imaging was obtained with a 2D PGSE sequence with slice direction along z, 300 *µ*m isotropic resolution, acquired with 38 slices and a matrix-size of 170×87, single line read-out, bandwidth = 18.5 kHz, TE/TR = 36/3200 ms. PGSE parameters were *δ* = 7.2 ms, Δ = 20.2 ms, varying gradient strength, and *b* = (50, 1000, 3000, 4000) s/mm^2^ and *b* = (50, 1000, 3000, 4000, 8000, 12000, 20000) s/mm^2^ for scan-series 1 and 2 respectively. Diffusion weighting was repeated with 21 uniformly distributed gradient vectors over a half-sphere. The directional scheme was repeated with opposite polarity for later correction of cross-terms between external diffusion gradients, image slice gradients, and macroscopic internal gradients [5]. The effect on the cross-terms between the external diffusion gradients and image slice gradients is shown in Sup. Fig. 13. Across all scans, the SNR in CC at *b* = 50 s/mm^2^ was 163±11 prior to noise suppression and 701±127 after.

Prior to the scan session the rotation setup with the agar embedded tissue was placed in the RF coil at room temperature at least 5 hours prior to scanning for temperature stabilization. To reduce short-term instabilities (i.e. temperature variations in the magnet and motion effect due to setup handling and initial heating of the gradient hardware) a dummy dMRI scan of >2 hours was collected before the actual DWI scans. To reduce temperature variation of the tissue during scanning a constant airflow of room temperature was ensured around the sample.

#### 4.3.3 Image post-processing

First, Gibbs ringing effects and noise were suppressed for all dMRI scans using gibbs_removal() and mppca() from the DIPY library [58]. Then, all scans were co-registered to a slightly rotated version of one of the scans using aling() from the DIPY library. To minimize cross-term effects between imaging gradients, macroscopic background field inhomogeneities and the diffusion gradients cross-term correction [5] was applied (Sup. Fig. 13).

### 4.4 Analysis

#### 4.4.1 Diffusion tensor modelling (in silico and ex vivo)

To assess the anisotropy of the apparent diffusion, we fitted the widely applied diffusion tensor (DT) model [22][23]. Although the model is not capable of fully characterising the non-Gaussian diffusion in white matter, it does capture the principal anisotropy of the signal. The DT model was fitted with the DIPY library [58] to the intervals *b* = (50, 1000, 3000, 4000) s/mm^2^, *b* = (4000, 8000, 12000) s/mm^2^, and *b* = (8000, 12000, 20000) s/mm^2^. The mean diffusivity (MD, mean of all eigenvalues), the axial diffusivity (AD, first eigenvalue), the radial diffusivity (RD, mean of second and third eigenvalue), and the first eigenvector (**e_1_**) were extracted for analysis.

In silico, orientation-dependency trends were categorized by quantifying minimum and maximum deviations between the cases with and without the integration of internal gradients. Ex vivo, orientation dependency was quantified based on the categories observed in silico by fitting each of the models

A. *f* (*x*) = *p*_0_ (no orientation-dependency)
B. *f* (*x*) = *p*_0_ + *p*_1_ *·* cos(*x*)^2^ (largest effect at angle(**B_0_**, **ẑ**) = 90 deg)
C. *f* (*x*) = *p*_0_ + *p*_1_ *·* cos(2*x*)^2^ (largest effect at angle(**B_0_**, **ẑ**) = 45 deg)
D. *f* (*x*) = *p*_0_ + *p*_1_ *·* cos(*x − π/*2)^2^ (largest effect at angle(**B_0_**, **ẑ**) = 0 deg)
 where *p*_0_ is a constant off-set, and *p*_1_ ≥ 0 represents the degree of orientation-dependency. The relative performances of the models are ranked by their respective Akaike information criterion (AIC) [59][60]

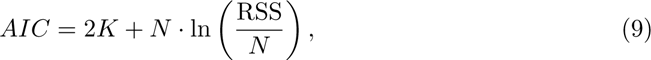

where *K* is the number of fitting parameters, *N* is the number of data points, and *RSS* is the residual sum of squares. To simplify the interpretation, the AIC-values were rescaled as

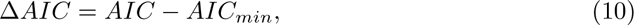

where *AIC_min_* is the minimum AIC obtained for any of the models. Δ*AIC* is the information loss experienced by fitting the given model rather than the best model. As suggested in [60] the relative support of the models is assessed according to the following scheme: Δ*AIC ≤* 2 indicates substantial support, 4 ≤ Δ*AIC ≤* 7 indicates considerably less support, and Δ*AIC >* 10 indicates essentially no support.

#### 4.4.2 Regions of interest from ex vivo images

We focus the analysis on the corpus callosum (CC) and the cingula (CING-L and CING-R). These bundles are oriented in the 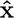 and **ŷ** directions of the field-of-view, respectively. With **ẑ** as the axis of rotation, CC will be parallel with **B_0_** when CING-L and CING-R are orthogonal with **B_0_**, and vice versa.

Voxels of interest were extracted based on manual segmentation in FSLeyes [61], and thresholding criteria: FA > 0.6, and angle with respect to the estimated fiber-direction of interest < 30 degrees.

## 5 Acknowledgements

S.W. has been supported by a DTU Alliance Stipend [project number 102682]. H.L. has received funding from the European Research Council (ERC) under the European Union’s Horizon 2020 research and innovation programme [grant agreement No 804746]. M.A. has been supported by Capital Region Research Foundation Grant [grant number A5657; principal investigator: T.B.D.]. T.B.D. has received funding from the European Research Council (ERC) under the European Union’s Horizon Europe research and innovation programme [grant agreement No 101044180].

## 6 Contributions

S.W., T.B.D. and H.L. designed the research. S.W. set up the experiments, wrote the code, acquired the data, and analyzed the data. S.W., T.B.D. and H.L. interpreted the results. T.B.D. provided tissue materials. M.A. and J.R-P. provided the XNH-informed meshes. S.W. wrote the majority of the paper. T.B.D., H.L., and M.A. contributed to writing the paper. T.B.D., H.L., M.A., J.R-P., and J-P.T. revised the paper.

## 7 Supplementary

**Figure 8:**
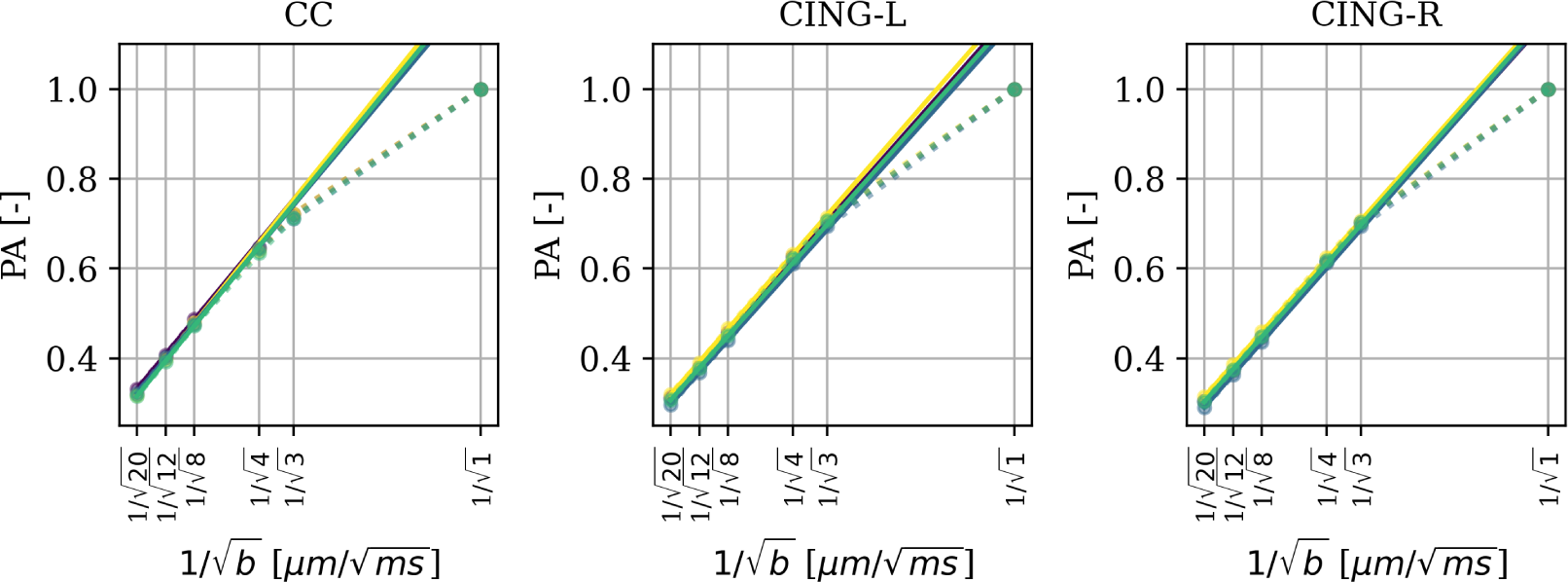
Validation of the signal in our high b-value regime (b=[8, 12, 20] ms/*µ*m^2^) expressing an isolated intra-axonal compartment by the powder average (PA) following a 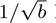 law [62][63]. Solid lines are linear fits over b=[8, 12, 20] ms/*µ*m^2^. With R2 *≥* 0.999 for all fits.

**Figure 9:**
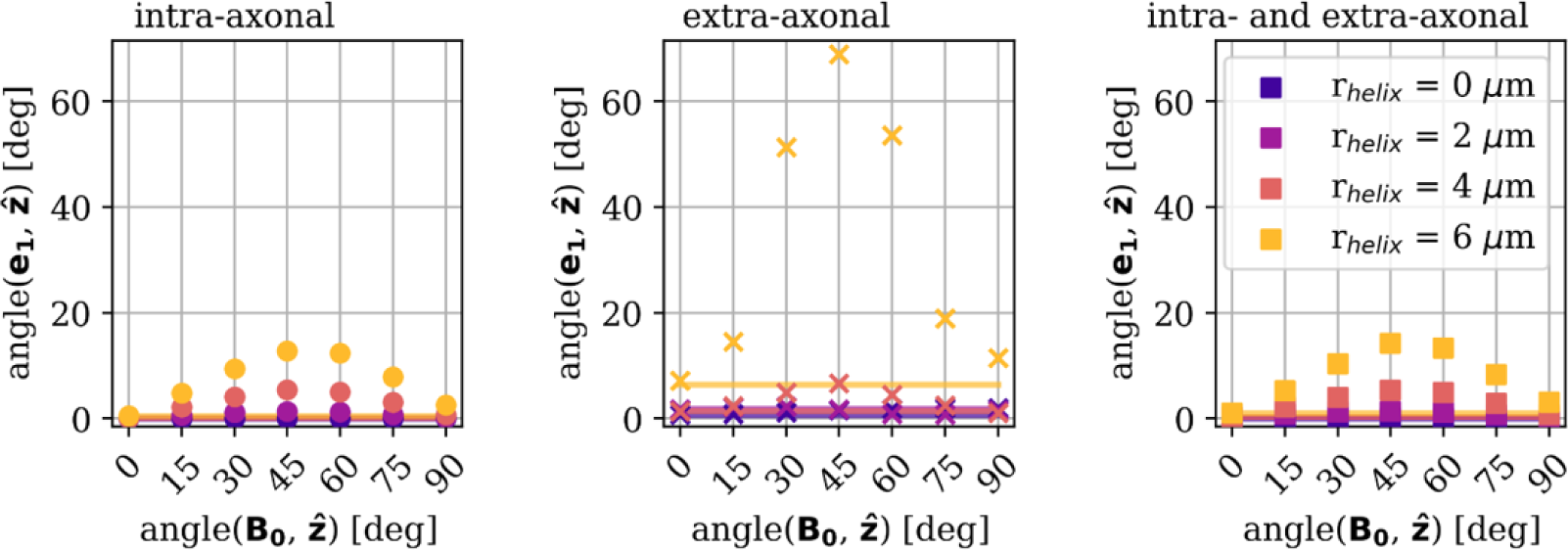
Susceptibility-induced internal fields affect the estimated fiber direction. We estimate the effect by computing the angle between the first eigenvector of the DT fit (**e_1_**) and the primary axis of the axons (**ẑ**). Markers indicate the DT metrics fitted to *S_eff_* (*b*). Lines indicate the DT metrics fitted to *S_des_*(*b*). It is seen that the angle is affected both by the helical radius of the axons *r_helix_* (i.e. the degree of undulation), and by *angle*(**ẑ***, B*_0_) (i.e. orientation w.r.t. **B_0_**). When no internal gradients are taken into account (lines) there is a substantial deviation between **ẑ** and **e_1_** for *r_helix_ ≥* 3.0*µ*m, as expected for high degrees of undulation. When internal gradients are taken into account (markers) the deviation between **e_1_** and **ẑ** becomes dependent on *angle*(**ẑ***, B*_0_) for *r_helix_ ≥* 4.0*µ*m. This orientation-dependency means that the estimated ADs and RDs will not be in the true axial and radial directions of the substrates, but instead biased by the susceptibility effects.

**Figure 10:**
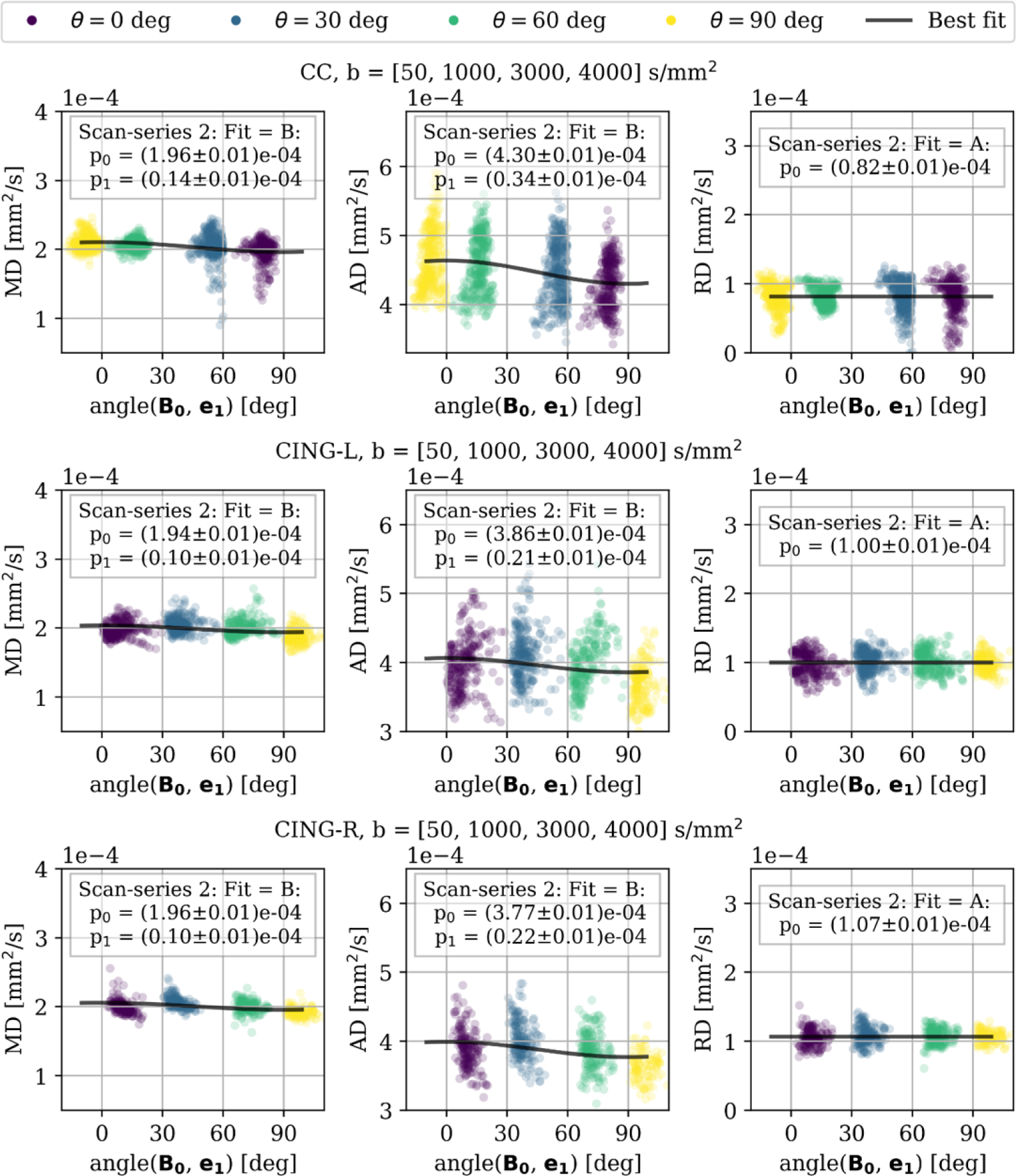
Scan-series 2. DT metrics fitted at low b-values ([50, 1000, 3000, 4000] s/mm^2^) for CC (upper row), CING-L (middle row), and CING-R (lower row) of an ex vivo monkey brain at 7 T. Markers are coloured according to which orientation the scan was acquired at, and plotted along with the best fit based on lowest AIC-value. Orientation-dependency is observed for AD of all ROIs, and for MD of CC and CING-L. The degree of orientation-dependency is stronger for CC than for CING-L and CING-R. The difference in effect on AD vs. RD shows that the PGSE signal is affected anisotropically. ΔAIC and ΔRMSE for all DT metrics and all fitted models are listed in App. Tab. 1

**Figure 11:**
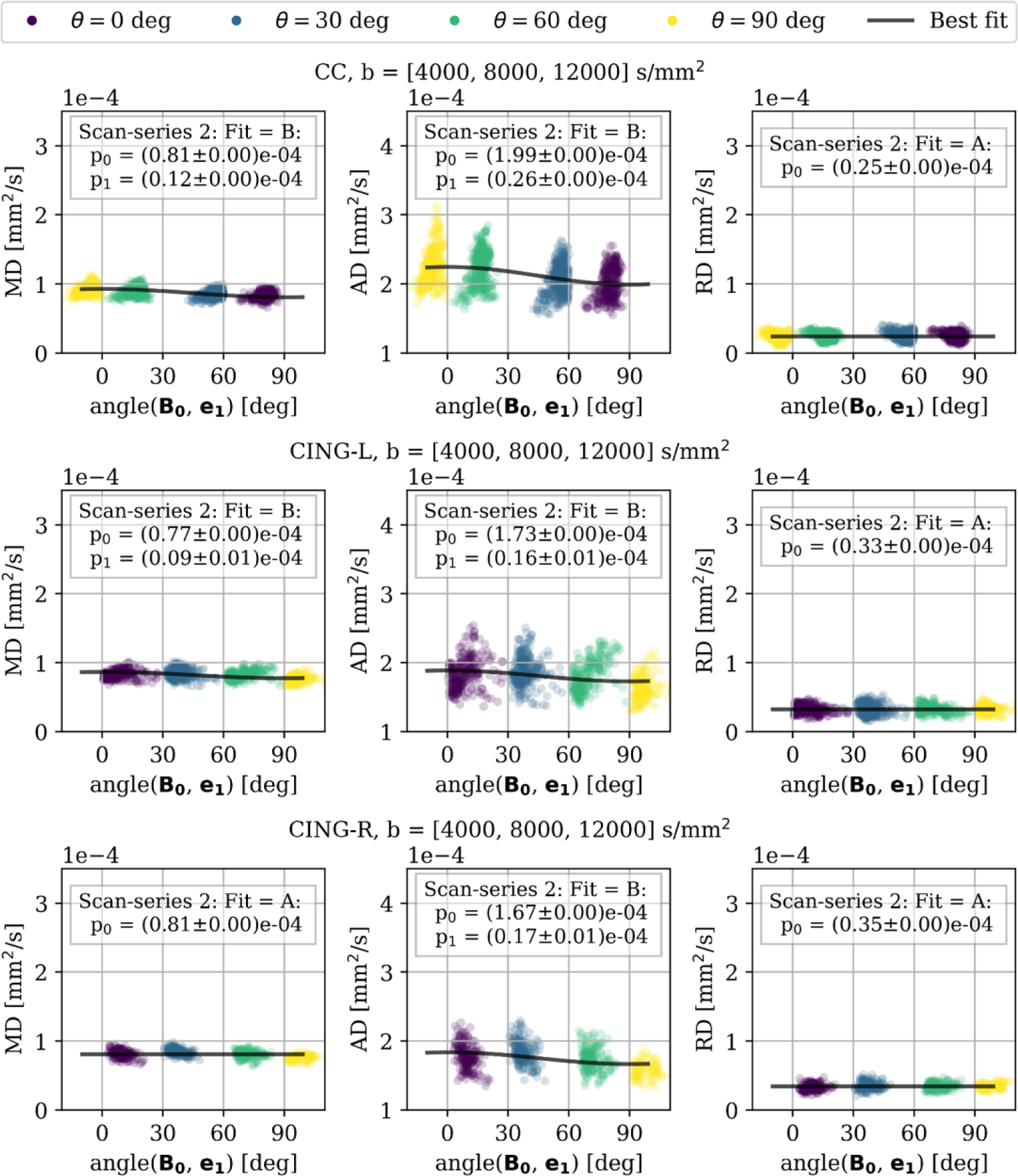
Scan-series 2. DT metrics fitted at intermediate b-values ([4000, 8000, 12000] s/mm^2^) for CC (upper row), CING-L (middle row), and CING-R (lower row) of an ex vivo monkey brain at 7 T. Markers are coloured according to which orientation the scan was acquired at, and plotted along with the best fit based on lowest AIC-value. Orientation-dependency is observed for AD of all ROIs, and for MD of CC. The degree of orientation-dependency is stronger for CC than for CING-L and CING-R. The difference in effect on AD vs. RD shows that the PGSE signal is affected anisotropically. ΔAIC and ΔRMSE for all DT metrics and all fitted models are listed in App. Tab. 2

**Figure 12:**
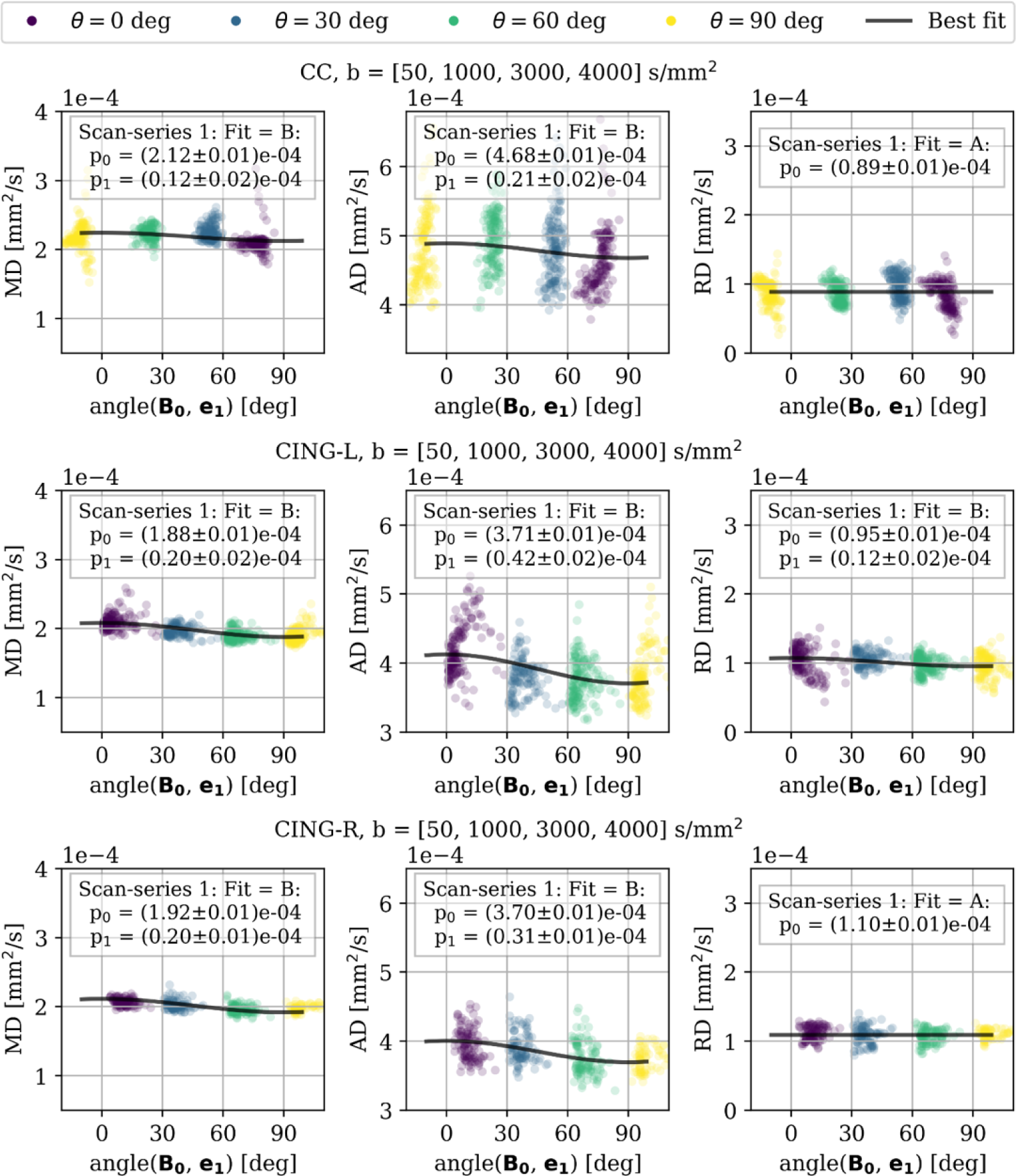
Scan-series 1. DT metrics fitted at low b-values ([50, 1000, 3000, 4000] s/mm^2^) for CC (upper row), CING-L (middle row), and CING-R (lower row) of an ex vivo monkey brain at 7 T. Markers are coloured according to which orientation the scan was acquired at, and plotted along with the best fit based on lowest AIC-value. Orientation-dependency is observed for AD of all ROIs, and for MD of CC. The degree of orientation-dependency is stronger for CC than for CING-L and CING-R. The difference in effect on AD vs. RD shows that the PGSE signal is affected anisotropically. ΔAIC and ΔRMSE for all DT metrics and all fitted models are listed in App. Tab. 4

**Figure 13:**
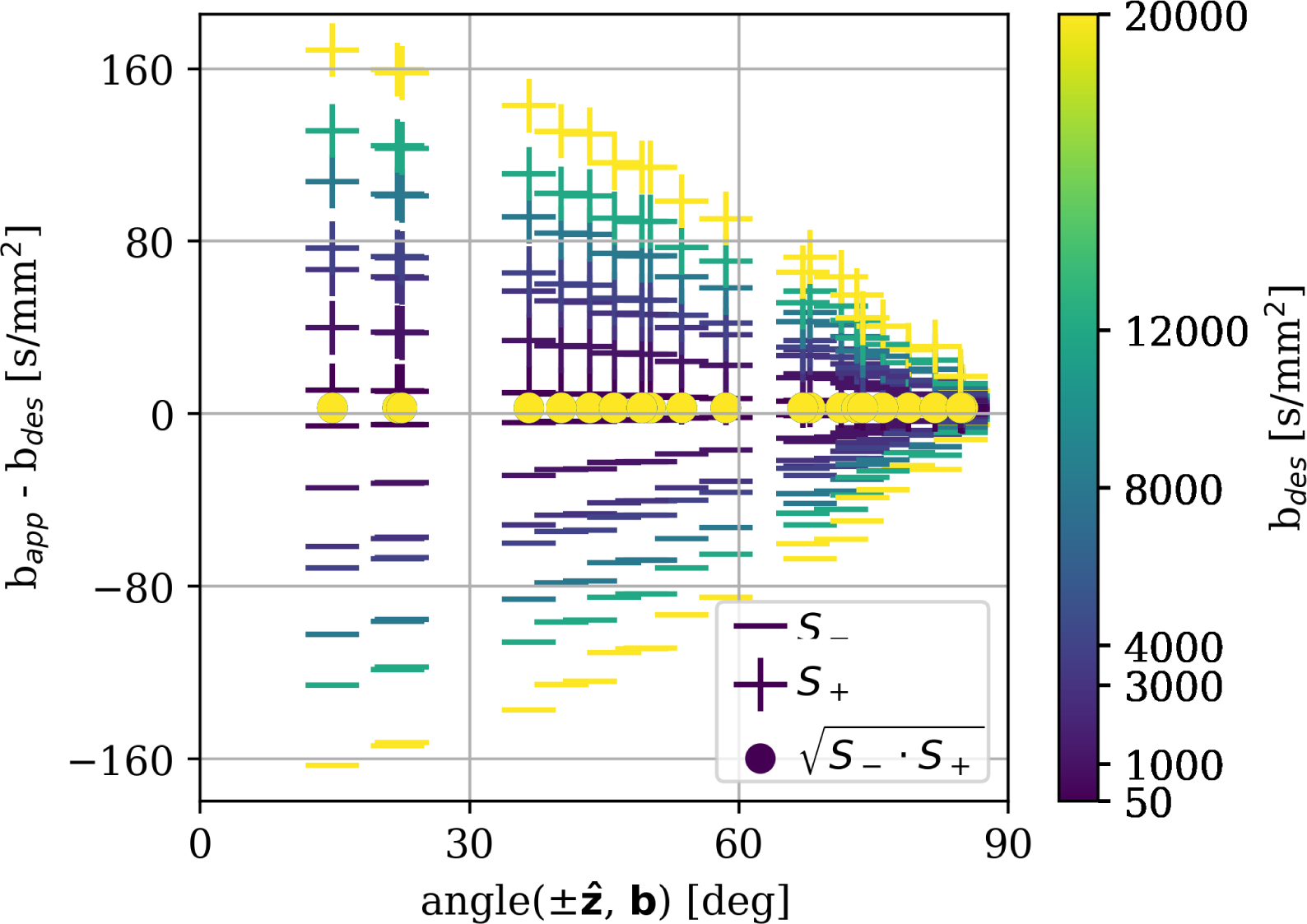
Cross-term correction of the applied b-values. For 2D imaging the applied b-values become skewed along the image-slice direction due to the super-position of slice gradients and diffusion gradients. +’s indicate the deviation between the applied b-value and the desired b-value for b-vectors with a positive component along **ẑ**. Vice versa for −’s. indicate the cross-term corrected b-value. For increasing b-values we see increasing anisotropy for the uncorrected values (i.e. the deviation of the magnitude of b-vectors at different orientations increases).

**Table 1:** Fitting performance according to ΔAIC for orientation-dependence of DT metrics of scan-series 2 at low b-values ([50, 1000, 3000, 4000] s/mm^2^) for CC (upper row), CING-L (middle row), and CING-R (lower row).

| $\Delta\text{AIC}$ for CC | | | |
| --- | --- | --- | --- |
| Model\Metric | MD | AD | RD |
| A | 143 | 116 | **0** |
| B | **0** | **0** | 8 |
| C | 112 | 112 | 12 |
| D | 151 | 119 | 22 |

| $\Delta\text{AIC}$ for CING-L | | | |
| --- | --- | --- | --- |
| Model\Metric | MD | AD | RD |
| A | 17 | 32 | **0** |
| B | **0** | **0** | 28 |
| C | 89 | 39 | 44 |
| D | 80 | 39 | 26 |

**Table 1:** Fitting performance according to $\Delta\text{AIC}$ for orientation-dependence of DT metrics of scan-series 2 at low b-values ([50, 1000, 3000, 4000] s/mm<sup>2</sup>) for CC (upper row), CING-L (middle row), and CING-R (lower row).
| $\Delta\text{AIC}$ for CING-R | | | |
| --- | --- | --- | --- |
| Model\Metric | MD | AD | RD |
| A | 21 | 50 | **0** |
| B | **0** | **0** | 36 |
| C | 98 | 57 | 46 |
| D | 93 | 57 | 27 |

**Table 2:** Fitting performance according to ΔAIC for orientation-dependence of DT metrics of scan-series 2 at intermediate b-values ([4000, 8000, 12000] s/mm^2^) for CC (upper row), CING-L (middle row), and CING-R (lower row).

| $\Delta\text{AIC}$ for CC | | | |
| --- | --- | --- | --- |
| Model\Metric | MD | AD | RD |
| A | 204 | 181 | **0** |
| B | **0** | **0** | 95 |
| C | 197 | 149 | 106 |
| D | 437 | 190 | 210 |

| $\Delta\text{AIC}$ for CING-L | | | |
| --- | --- | --- | --- |
| Model\Metric | MD | AD | RD |
| A | 7 | 49 | **0** |
| B | **0** | **0** | 124 |
| C | 200 | 71 | 163 |
| D | 183 | 67 | 145 |

**Table 2:** Fitting performance according to $\Delta\text{AIC}$ for orientation-dependence of DT metrics of scan-series 2 at intermediate b-values ([4000, 8000, 12000] s/mm<sup>2</sup>) for CC (upper row), CING-L (middle row), and CING-R (lower row).
| $\Delta\text{AIC}$ for CING-R | | | |
| --- | --- | --- | --- |
| Model\Metric | MD | AD | RD |
| A | **0** | 67 | **0** |
| B | 1 | **0** | 162 |
| C | 189 | 89 | 212 |
| D | 170 | 86 | 141 |

**Table 3:** Fitting performance according to ΔAIC for orientation-dependence of DT metrics of scan-series 2 at high b-values ([8000, 12000, 20000] s/mm^2^) for CC (upper row), CING-L (middle row), and CING-R (lower row).

| $\Delta\text{AIC}$ for CC | | | |
| --- | --- | --- | --- |
| Model\Metric | MD | AD | RD |
| A | 280 | 212 | **0** |
| B | **0** | **0** | 327 |
| C | 252 | 151 | 403 |
| D | 622 | 237 | 895 |

| $\Delta\text{AIC}$ for CING-L | | | |
| --- | --- | --- | --- |
| Model\Metric | MD | AD | RD |
| A | **0** | 33 | **0** |
| B | 126 | **0** | 594 |
| C | 363 | 79 | 734 |
| D | 323 | 74 | 601 |

**Table 3:** Fitting performance according to $\Delta\text{AIC}$ for orientation-dependence of DT metrics of scan-series 2 at high b-values ([8000, 12000, 20000] s/mm<sup>2</sup>) for CC (upper row), CING-L (middle row), and CING-R (lower row).
| $\Delta\text{AIC}$ for CING-R | | | |
| --- | --- | --- | --- |
| Model\Metric | MD | AD | RD |
| A | **0** | 71 | **0** |
| B | 149 | **0** | 436 |
| C | 386 | 122 | 502 |
| D | 341 | 116 | 322 |

**Table 4:** Fitting performance according to ΔAIC for orientation-dependence of DT metrics of scan-series 1 at low b-values ([50, 1000, 3000, 4000] s/mm^2^) for CC (upper row), CING-L (middle row), and CING-R (lower row).

| $\Delta\text{AIC}$ for CC | | | | |
| --- | --- | --- | --- | --- |
| Model\Metric | MD | AD | RD |  |
| A | 11 | 12 | **0** |  |
| B | **0** | **0** | 12 |  |
| C | 34 | 16 | 16 |  |
| D | 33 | 15 | 15 |  |

| $\Delta\text{AIC}$ for CING-L | | | | |
| --- | --- | --- | --- | --- |
| Model\Metric | MD | AD | RD |  |
| A | 284 | 111 | 27 |  |
| B | **0** | **0** | **0** |  |
| C | 286 | 87 | 55 |  |
| D | 320 | 114 | 59 |  |

**Table 4:** Fitting performance according to $\Delta\text{AIC}$ for orientation-dependence of DT metrics of scan-series 1 at low b-values ([50, 1000, 3000, 4000] s/mm<sup>2</sup>) for CC (upper row), CING-L (middle row), and CING-R (lower row).
| $\Delta\text{AIC}$ for CING-R | | | | |
| --- | --- | --- | --- | --- |
| Model\Metric | MD | AD | RD |  |
| A | 64 | 75 | **0** |  |
| B | **0** | **0** | 60 |  |
| C | 108 | 84 | 13 |  |
| D | 133 | 102 | 35 |  |

## Notes

### Competing Interest Statement

The authors have declared no competing interest.

